# Sex peptide-SPR-serotonin signaling regulates mating-induced Sex peptide replenishment in *Drosophila*, suggesting an ancestral role in males

**DOI:** 10.64898/2026.10.19.760009

**Authors:** Ryo Hoshino, Yasuhiro Kozono, Ryoya Tanaka, Hsin-Kuang Lin, Tomotsune Ameku, Tomohiro Tei, Tsunaki Asano, Azusa Kamikouchi, Yuko Shimada-Niwa, Akira Nakamura, Takeshi Awasaki, Ryusuke Niwa

## Abstract

Seminal fluid not only assists successful fertilization but also induces changes in female post-mating behavior and physiology. Consequently, males with depleted or inadequate seminal fluid produce fewer offspring. Males of iteroparous species therefore increase seminal fluid production to replenish exhausted supplies. However, it remains unclear how mating-induced replenishment of seminal fluid components is regulated at the molecular and cellular levels. Here, we show that replenishment of Sex peptide (SP) in male *Drosophila melanogaster* requires both SP and its receptor SPR. Mechanistically, mating-induced SP replenishment in the male accessory gland (MAG) is regulated by MAG-projecting serotonergic neurons, and this regulation depends on SPR, which is likely activated by SP as it passes through the reproductive tract during mating. This inter-tissue signaling pathway is not limited to SP but also coordinates the post-mating upregulation of an additional seminal fluid protein gene. While SP-SPR signaling is a well-known regulator of female post-mating responses, we demonstrate that it also modulates male physiology. Furthermore, comparative analyses across 12 drosophilid species are consistent with the hypothesis that SP-SPR signaling acquired a role in males before its role in canonical female post-mating responses evolved. We therefore propose that the role of SP-SPR signaling as a mating signal in males is ancestral to its more familiar but evolutionarily derived role in females.

## Introduction

Reproductive success requires the transfer of seminal fluid from males to females during mating. Seminal fluid comprises both spermatozoa and seminal plasma, the non-sperm components. Seminal plasma is a complex cocktail of lipids, proteins, carbohydrates, salts, hormones, nucleic acids, vitamins, and other molecules that profoundly influence reproductive success because it is important for sperm stability and delivery (Poiani, 2006; Scolari et al., 2021). Moreover, its components can also modulate female physiology, behavior, and reproductive output after mating. Collectively, these mating-induced changes are called post-mating responses (PMRs) (Avila et al., 2011; Hopkins and Perry, 2022; Poiani, 2006). From insects to mammals, seminal fluid has evolved as a key mediator of sexual communication and conflict, acting at the interface between individuals to shape reproductive strategies and outcomes (Poiani, 2006). The seminal fluid of malaria mosquitoes, for example, contains male-derived ecdysteroids that activate oogenesis and suppress female sexual receptivity (Peng et al., 2022; Pondeville et al., 2008). In induced ovulators, such as rabbits, alpacas, and llamas, seminal plasma triggers ovulation via nerve growth factor signaling (Bakker and Baum, 2000; Maranesi et al., 2018). In placental mammals, including humans, cytokines and prostaglandins in seminal fluid modulate the female immune response (Bromfield, 2014; Schjenken and Robertson, 2014).

In the fruit fly *Drosophila melanogaster*, Sex peptide (SP), a 36-amino acid seminal fluid peptide produced in the male accessory gland (MAG), acts through SP receptor (SPR) expressed in female SP-sensing sensory neurons (f-SPSNs) (Chapman et al., 2003; Chen et al., 1988; Feng et al., 2014; Häsemeyer et al., 2009; Heifetz et al., 2001; Hopkins and Perry, 2022; Liu and Kubli, 2003; Qazi et al., 2003; Yapici et al., 2008). SP–SPR signaling is essential for a wide variety of PMRs, including changes in mating rejection, feeding, egg laying, immunity, and oogenesis (Hoshino and Niwa, 2021; Kubli, 2003). Consistent with this central role, loss or insufficiency of SP markedly alters female PMRs. Accordingly, the transfer of insufficient SP results in reduced offspring production (Liu and Kubli, 2003). In addition, the lack of SP-induced suppression of remating further reduces male reproductive fitness because of the resulting increase in sperm competition (Misra and Wolfner, 2020). Males also transfer progressively less SP during successive matings (Hihara, 1981; Sirot et al., 2009), but can subsequently increase SP production to replenish depleted stores (Hollis et al., 2019; Koppik et al., 2018; Sirot et al., 2009). Although mating activates expression of several seminal fluid protein genes, how males sense mating-associated SP release and how they regulate its replenishment remain unknown (Bertram et al., 1992; DiBenedetto et al., 1990; Herndon et al., 1997; Monsma et al., 1990; Schmidt et al., 1985; Simmerl et al., 1995).

Here, we report that the replenishment of SP in *D. melanogaster* males requires both SP and SPR. Mechanistically, mating-induced replenishment in the MAG is regulated by serotonin released from MAG-projecting serotonergic neurons, and their activity depends on SPR. To the best of our knowledge, ours is the first study to demonstrate that SP–SPR signaling also controls male physiology. Moreover, in contrast to the role of SP–SPR signaling in female PMRs (Kubli and Bopp, 2012), we show conservation of the male SP replenishment mechanism even to Sophophora species outside of the *melanogaster* and *ananassae* groups. We propose that the role of SP–SPR signaling as a mating signal in males predates its role in triggering female PMRs.

## Results

### Mating experience induces SP replenishment and *SP* transcription

To determine the precise time required for SP replenishment after mating, we performed a capillary western blotting assay to measure temporal changes in the amount of SP protein stored in the male reproductive tract. The male mating paradigm in this study is illustrated in Fig. 1A. First, we collected young males that had never mated (“naive” hereafter) 5 days after eclosion and placed them in vials containing standard food and virgin females at a ratio of 1 male to 3 females. After 24 hrs of multiple matings, wild-type males that had mated with females (“experienced” hereafter) retained approximately 10% of their initial SP protein levels (Fig. 1B; compare lanes 1 and 2; also see Fig. 1C and 1D). We then isolated the newly experienced males in new vials and quantified SP in their reproductive tracts after 24, 48, 72, and 96 hrs. Although wild-type males replenished SP by approximately 40% within 24 hrs, its production slowed down thereafter (24–96 hrs) (Fig. 1B; lanes 3 to 10; also see Fig. 1C and 1D). These results suggest that SP replenishment is actively induced within 0–24 hrs post-mating. For this reason, hereafter, we mainly focused on the replenishment process in the 24 hrs after mating.

**Figure 1.**
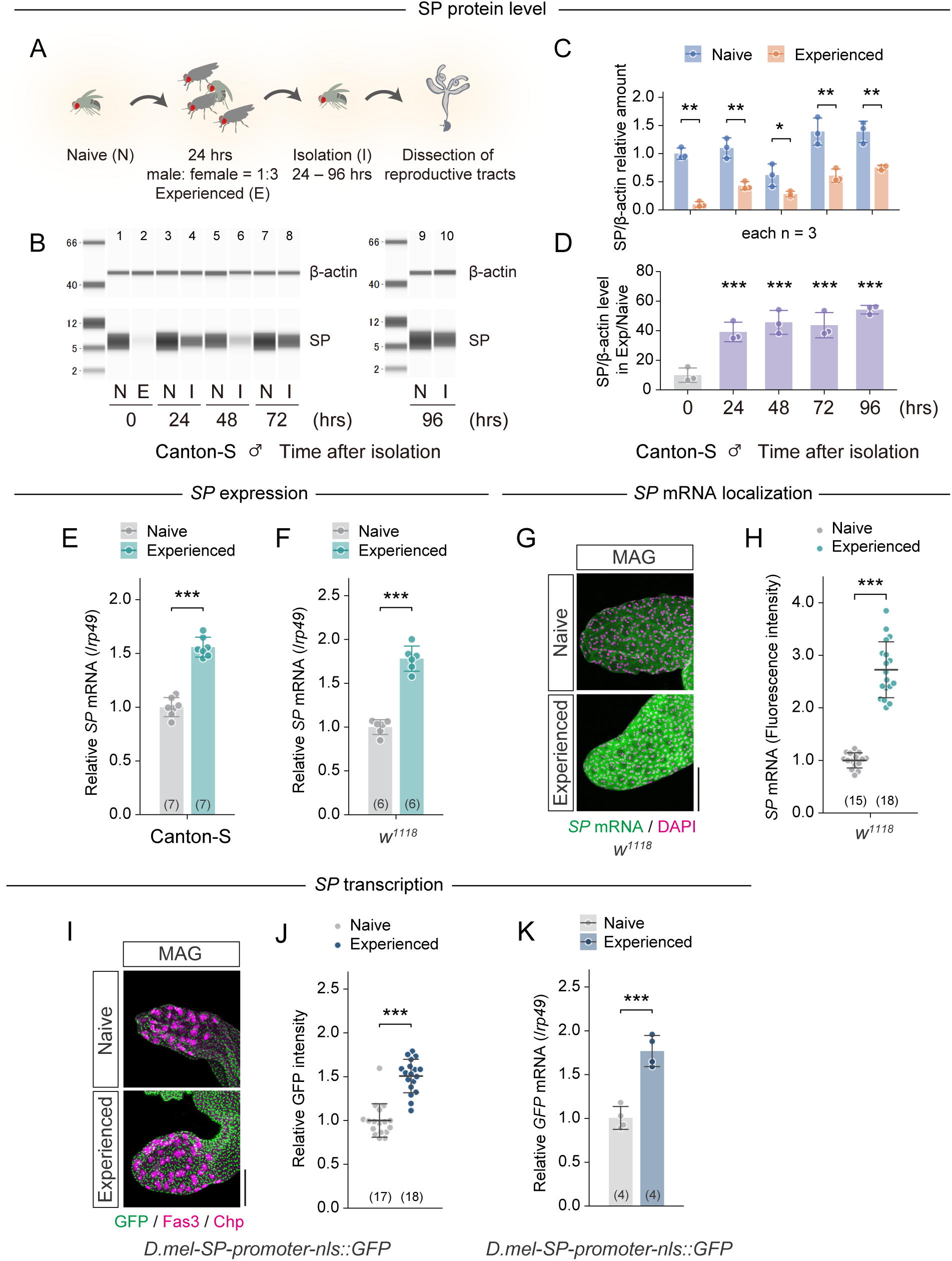
Mating promotes SP replenishment via transcriptional regulation. (A–D) Temporal changes in SP protein in the Canton-S male reproductive tract following mating. (A) Schematic showing the mating protocol followed by dissection and SP quantification. (B) Representative images of western blots using anti-SP (7 kDa) and anti-β-actin (47 kDa) antibodies. Lane numbers indicated above each lane. (C) Quantification of SP protein levels in the male reproductive tract normalized to β-actin. (D) Temporal changes in SP protein levels in experienced males compared to naive males. Statistical significance was determined by comparing each sample to the 0 hr sample. (E–F) *SP* mRNA levels in male (E) Canton-S and (F) *w^1118^* reproductive tracts before and after mating measured via RT-qPCR. *SP* mRNA levels were calculated using *rp49* as the reference gene and subsequently normalized to the mean level in naive males. (G–H) *SP* mRNA levels in *w^1118^* MAG before and after mating as visualized by *in situ* HCR. (G) Representative images of *SP* mRNA (green) and DAPI (magenta) staining at the tip of the MAG. Scale bar, 100 μm. (H) Quantification of *SP* mRNA signal intensity in the MAG. Signal intensities were normalized to the mean signal intensity of naive males. (I–K) Quantification of *SP* promoter-nls::GFP reporter activity before and after mating. (I) Representative images of MAGs immunostained with anti-GFP (green), anti-Fas3 (main cell marker; magenta), and anti-Chp (secondary cell marker; magenta) antibodies. Scale bar, 100 μm. Note that the most prominent magenta signals in the apical side of the MAG correspond to secondary cells, which are anti-Chp-positive. (J) Quantification of nls::GFP intensity levels in the MAGs of *w^1118^* males. Signal intensities were normalized to the mean signal intensity of naive males. (K) *GFP* mRNA levels in male reproductive tracts before and after mating measured via RT-qPCR. *GFP* mRNA levels were calculated using *rp49* as the reference gene and subsequently normalized to the mean level in naive males. All quantitative data are presented as means ± SD. The number of samples (n) is indicated in each graph. Statistical analysis: unpaired two-tailed Student’s *t*-test for (C, E, F, and K); Dunnett’s test for (D); and Wilcoxon rank sum test for (H and J). \**P* < 0.05, \*\**P* < 0.01, \*\*\**P* < 0.001; n.s., not significant (*P* ≧ 0.05).

Several studies showed that mating activates the transcription of seminal fluid protein genes in the male reproductive tract (Bertram et al., 1992; DiBenedetto et al., 1990; Herndon et al., 1997), whereas post-mating induction of *SP* transcripts has not been examined. We observed via RT-qPCR and *in situ* hybridization chain reaction (isHCR) an increase in *SP* mRNA levels in wild-type and control males 24 hrs after mating (Fig. 1, E–H). This post-mating increase in *SP* mRNA in the MAG appears to be transcriptionally regulated, because we observed a similar mating-induced increase in the expression of a GFP reporter driven by a genomic fragment containing a putative *SP* promoter/enhancer region (∼1.6 kbp upstream from the *SP* start codon) (Fig. 1, I–K). These results suggest that post-mating SP replenishment involves increased *SP* transcription.

### The post-mating increase in *SP* mRNA is triggered by ejaculation

Corazonin (Crz)-producing neurons control ejaculation (Tayler et al., 2012). We found that inhibiting ejaculation by silencing Crz neurons or Crz receptor (Crz-R) neurons suppressed the mating-induced increase in *SP* transcription (Supplementary Fig. S1, A and B). Conversely, artificially inducing ejaculation by activating Crz neurons increased *SP* mRNA even in the absence of mating (Supplementary Fig. S1C). These results imply that ejaculation triggers the post-mating induction of *SP*. However, *Crz* and *Crz-R* loss-of-function mutant males, which can ejaculate normally and are fertile (Khan et al., 2021; Sha, 2013; Sha et al., 2014), showed mating-induced increases in *SP* mRNA similar to those of control males (Supplementary Fig. S1, D and E). These results suggest that although ejaculation influences the mating-induced *SP* mRNA increase, it does so independently of Crz–Crz-R signaling.

### Neuronal serotonin is required for the post-mating increase in mRNAs of *SP* and some other accessory gland protein genes

We next investigated the signal acting on the MAG to regulate post-mating SP replenishment. We focused on serotonin because serotonergic neurons in the posterior ventral nerve cord (VNC) project directly to the MAG (Jois et al., 2018), and serotonin injection can evoke sperm ejection (Tayler et al., 2012).

We first used the driver *Tryptophan hydroxylase neuronal* (*Trhn*)-*GAL4*, which labels serotonergic neurons, to determine whether manipulating these neurons regulates *SP* mRNA levels. Silencing serotonergic neurons prevented the post-mating increase in *SP* mRNA, whereas their artificial activation increased *SP* mRNA in naive males (Supplementary Fig. S2, A and B). These results suggest that serotonergic neuronal activity affects *SP* transcription.

We next examined the requirement of serotonin biosynthesis. *De novo* serotonin biosynthesis requires the conversion of tryptophan to 5-hydroxytryptophan (5-HTP) (Fig. 2A). In *D. melanogaster*, this conversion occurs both in neurons and peripheral non-neuronal tissues in reactions mediated by Trhn and Henna, respectively (Rosikon et al., 2023). We therefore generated Trhn and Henna null mutants using CRISPR-Cas9 technology (Supplementary Fig. S2, C and D). *Trhn*-null males showed a significantly blunted post-mating increase in *SP* at the mRNA level (Fig. 2, B and C) accompanied by a trend toward a reduction in SP protein replenishment (Fig. 2D; compare lane 3 with lane 6; also see Fig. 2E). In contrast, we still observed the post-mating increase in *SP* mRNA in *Henna*-null males (Supplementary Fig. S2E). This suggests that neuronal serotonin is essential for SP replenishment after mating.

**Figure 2.**
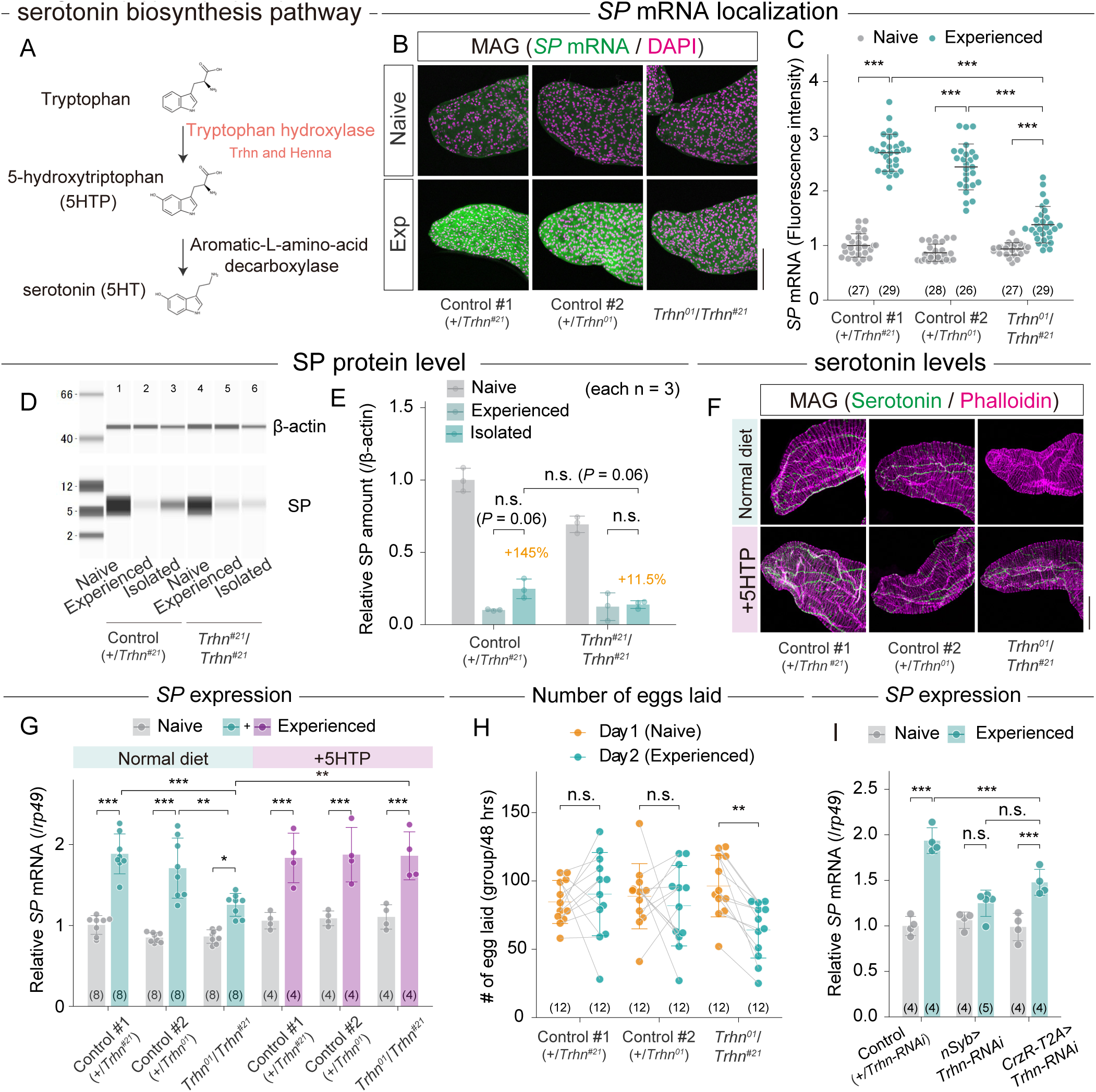
MAG-projecting serotonergic neurons facilitate *SP* transcription. (A) The serotonin biosynthesis pathway. (B–C) *SP* mRNA levels before and after mating in the MAG of the *Trhn* null mutant (*Trhn^01^/Trhn^#21^*) and controls (*+/Trhn^01^*and *+/Trhn^#21^*) visualized via *in situ* HCR. (B) Representative images of *SP* mRNA (green) and DAPI (magenta) staining in the MAG. Scale bar, 100 μm. (C) Quantification of *SP* mRNA intensity in the MAG. Signal intensities were normalized to the mean signal intensity of control males (*+/Trhn^01^*). (D-E) Temporal changes in SP protein levels before and after mating in *Trhn* null mutant (*Trhn^#21^/Trhn^#21^*) and control (*+/Trhn^#21^*) male reproductive tracts. (D) Representative images of western blots using anti-SP (7 kDa) and anti-β-actin (47 kDa) antibodies. Lane numbers are indicated above each lane. (E) Quantification of SP protein levels normalized to β-actin. Values are shown relative to the mean of the naive control males (*+/Trhn^#21^*). (F) Representative images of anti-serotonin immunostaining (green) and phalloidin staining (magenta) in the male reproductive tracts of *Trhn* null mutant (*Trhn^01^/Trhn^#21^*) and controls (*+/Trhn^#21^*and *+/Trhn^01^*) males fed a standard diet with or without 5-HTP. Scale bar, 100 μm. (G) *SP* mRNA levels before and after mating in the male reproductive tract of *Trhn* null mutant (*Trhn^01^/Trhn^#21^*) and controls (*+/Trhn^#21^* and *+/Trhn^01^*) males, as measured by RT-qPCR. *SP* mRNA levels were calculated using *rp49* as the reference gene and subsequently normalized to the mean level in naive control males (*+/Trhn^01^*). Males were fed a standard diet with or without 5-HTP for 4 days until dissection. (H) Eggs laid by females mated to *Trhn* null mutant (*Trhn^01^/Trhn^#21^*) or controls (*+/Trhn^01^* and *+/Trhn^#21^*) males. See Materials and Methods for experimental details. For both Day-1 and Day-2 females, the number of laid eggs was counted over 48 hrs—the 24-hr male– female contact period followed by 24 hrs of isolation. (I) Pre- and post-mating *SP* mRNA levels in the male reproductive tract of flies carrying *nSyb-GAL4* or *CrzR-T2A-GAL4* and *Trhn*-RNAi transgenes, as measured via RT-qPCR. *SP* mRNA levels were calculated using *rp49* as the reference gene and subsequently normalized to the mean level in naive control males (+/*Trhn*-RNAi). All quantitative data are presented as means ± SD. The number of samples (n) is indicated in each graph. Statistical analysis: Steel-Dwass’s multiple comparison test for (C); unpaired two-tailed Student’s *t*-test for (E); Tukey-Kramer’s HSD test for (G and I); and paired two-tailed Student’s *t*-test for (H). \**P* < 0.05, \*\**P* < 0.01, \*\*\**P* < 0.001; n.s., not significant (*P* ≧ 0.05).

We speculated that oral administration of 5-HTP may restore serotonin synthesis in *Trhn*-null males and rescue the post-mating increase in *SP* mRNA. After confirming that *Trhn*-null males showed loss of anti-serotonin immunoreactivity in the axonal terminals of the MAG-projecting serotonergic neurons, we confirmed that oral 5-HTP feeding restored anti-serotonin immunoreactivity in *Trhn*-null males (Fig. 2F and Supplementary Fig. S2F). Importantly, 5-HTP administration also restored *SP* mRNA levels in experienced *Trhn*-null males while having no significant effect on naive *Trhn*-null males (Fig. 2G).

Because several genes encoding accessory gland proteins (ACPs) are transcriptionally upregulated after mating (Bertram et al., 1992; DiBenedetto et al., 1990; Herndon et al., 1997), we asked whether neuronal serotonin regulates ACP expression more broadly. RNA-seq of 155 annotated ACPs (McQuarrie et al., 2025) identified 47 with significant post-mating upregulation in controls and 35 in *Trhn*-null mutants (FC ≧ 1.5, adjusted *P* ≦ 0.05); 18 were upregulated only in controls (Supplementary Fig. S3, A and B). These results suggest that neuronal serotonin contributes to the post-mating transcriptional induction of a subset of ACP genes.

To test the reproductive consequences of impaired ACP replenishment, we sequentially exposed individual males to two sets of three virgin females. Females mated with *Trhn*-null males, but not with control males, showed reduced egg laying during the second mating round (Fig. 2H). GFP-labeled sperm confirmed that *Trhn*-null males efficiently inseminated all three females in most replicates (Supplementary Fig. S3, C and D), arguing against a complete failure of sperm transfer. Taken together with the transcriptional and protein-level data described above, these findings indicate that neuronal serotonin is required for post-mating SP replenishment.

### MAG-projecting serotonergic neurons in the posterior VNC are required for post-mating SP replenishment

To test the requirement for MAG-projecting serotonergic neurons, we knocked down *Trhn* using *CrzR-T2A-GAL4*, which labels a subset of posterior VNC serotonergic neurons projecting to the MAG (Tayler et al., 2012). This significantly reduced anti-serotonin immunoreactivity in their cell bodies and MAG projections (Supplementary Fig. S4, A–C). *Trhn* knockdown also suppressed the post-mating increase in *SP* mRNA, although the effect was weaker than what we observed with a pan-neuronal driver (Fig. 2I). These findings support the idea that the MAG-projecting serotonergic neurons evoke the mating-induced increase in *SP* mRNA.

### Serotonin administration is sufficient to induce *SP* mRNA transcription and enhance protein synthesis *ex vivo*

To test whether serotonin induces *SP* transcription, we first injected serotonin into naive males and observed increased *SP* mRNA after 3 hrs (Supplementary Fig. S5A). Serotonin also increased *SP* mRNA in *ex vivo*-cultured male reproductive tracts after 6–16 hrs (Fig. 3, A and B), with significant effects at concentrations ≥10 μM (Supplementary Fig. S5B).

**Figure 3.**
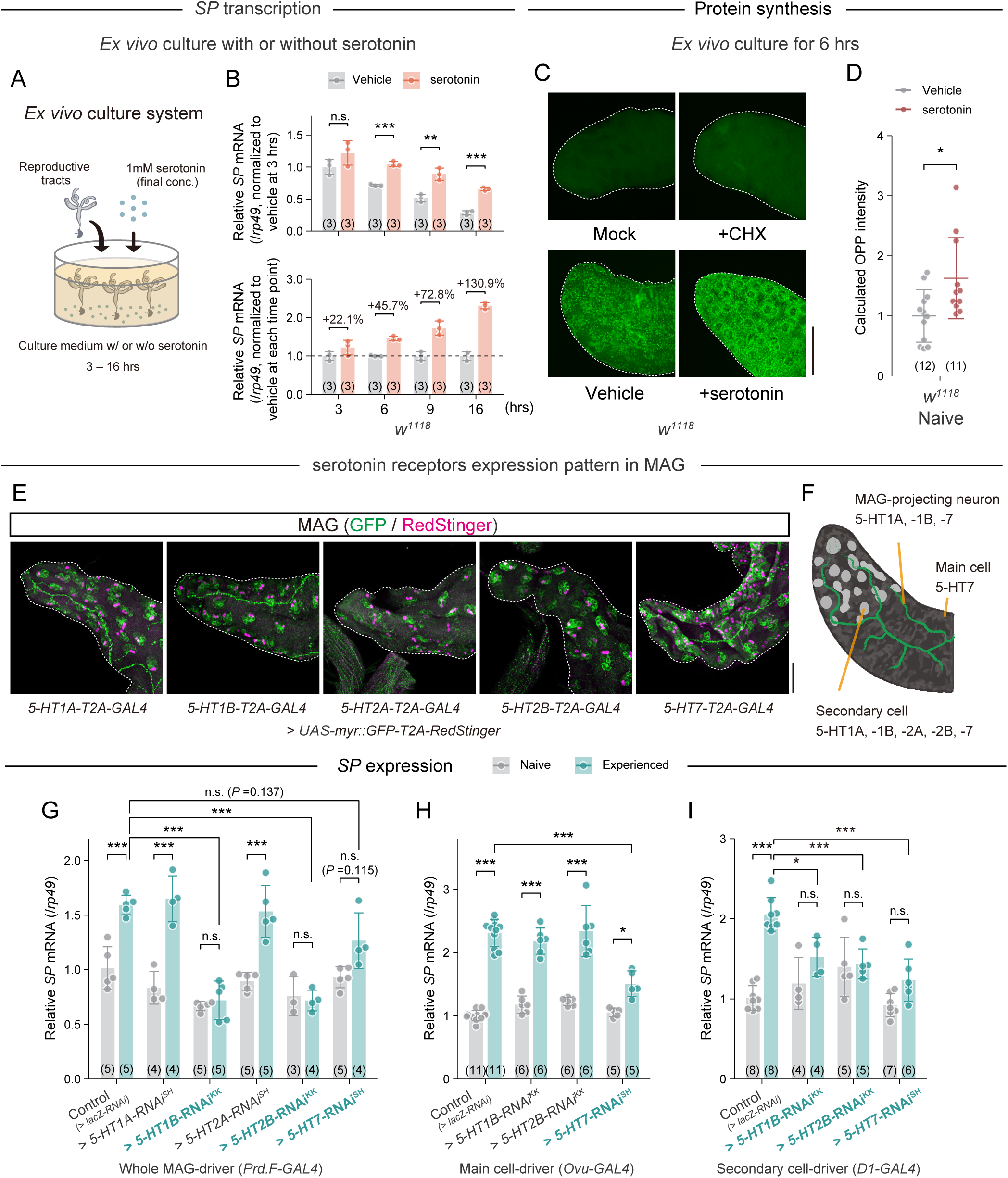
Serotonin reception in the MAG is required for mating-induced *SP* mRNA elevation. (A) Schematic for the *ex vivo* culture of male reproductive tracts. (B) *SP* mRNA levels in male reproductive tracts cultured *ex vivo* with or without 1 mM serotonin for 3, 6, 9, and 16 hrs, measured via RT-qPCR and normalized to *rp49*. The upper panel shows *SP* mRNA levels at various time points, normalized to the vehicle control at 3 hrs. The lower panel shows the fold change in the relative *SP* mRNA levels between the control and serotonin-treated samples for each time point. (C and D) *Ex vivo* OPP incorporation assay with MAGs from naive *w^1118^* males. (C) Representative images showing OPP signals (green) in MAGs (outlined by dashed lines) from naive males following *ex vivo* culture with vehicle, serotonin, or cycloheximide (CHX). Samples incubated without OPP served as a negative control (mock). Scale bar, 100 µm. (D) Quantification of OPP fluorescence intensity in *ex vivo*-cultured MAGs. Signal intensities were normalized to the mean signal intensity of the vehicle group. (E and F) Expression of serotonin receptor genes in the MAG. (E) Representative images of the MAG from five serotonin receptor-*T2A-GAL4* lines driving the membrane-bound myr::GFP (green) and the nuclear marker RedStinger (magenta). Scale bar, 50 µm. (F) Schematic of the distributions of serotonin receptors in the MAG. (G) *SP* mRNA levels before and after mating in males with *Prd.F-GAL4* driving serotonin receptor knockdown in the whole MAG, measured via RT-qPCR. *SP* mRNA levels were calculated using *rp49* as the reference gene and subsequently normalized to the mean level in control males (*Prd.F-GAL4* > *lacZ-RNAi*). (H and I) *SP* mRNA levels before and after mating in males *with Ovu-GAL4* (H) or *D1-GAL4* (I) driving knockdown of *5-HT1B*, *5-HT2B*, or *5-HT7* in main cells or secondary cells, respectively. *SP* mRNA levels were measured via RT-qPCR. *SP* mRNA levels were calculated using *rp49* as the reference gene and subsequently normalized to the mean level in control males [*Ovu-GAL4* > *lacZ-RNAi* (H) or *D1-GAL4* > *lacZ-RNAi* (I)]. All quantitative data are presented as means ± SD. The number of samples (n) is indicated in each graph. Statistical analysis: unpaired two-tailed Student’s *t*-test for (B); Wilcoxon rank sum test for (D); and Tukey-Kramer’s HSD test for (G, H, and I). \**P* < 0.05, \*\**P* < 0.01, \*\*\**P* < 0.001; n.s., not significant (*P* ≧ 0.05).

Given that ribosome biosynthesis in the MAG increases after mating (Schmidt et al., 1985), we also examined protein production using *O*-propargyl puromycin (OPP). Serotonin administration significantly increased OPP incorporation in cultured MAGs (Fig. 3, C and D), indicating enhanced protein production. Consistent with our RNA-seq data showing that serotonin is required for the post-mating upregulation of multiple ACPs, our results suggest that serotonin broadly enhances the replenishment of multiple ACPs, rather than acting exclusively on SP.

### Post-mating SP replenishment requires several serotonin receptor subtypes expressed in MAG cells

The *D. melanogaster* genome encodes five serotonin receptor subtypes: *5-HT1A*, *5-HT1B*, *5-HT2A*, *5-HT2B*, and *5-HT7* (Rosikon et al., 2023). Using receptor *T2A-GAL4* knock-in lines (Deng et al., 2019) crossed with fluorescent protein reporter lines, we detected expression of *5-HT7* in main cells, whereas expression of all five receptors was detected in secondary cells (Fig. 3, E and F).

We then performed RNAi-mediated knockdown of each receptor gene using the MAG driver *Prd.F-GAL4*, which targets both main and secondary cells (Sharma et al., 2017; Xue and Noll, 2002). Knockdown of *5-HT1B*, *5-HT2B*, or *5-HT7* but not *5-HT1A* or *5-HT2A* significantly suppressed *SP* mRNA elevation after mating (Fig. 3G). Consistent with this result, we did not observe the post-mating increase in *SP* mRNA in genetic loss-of-function mutants for *5-HT1B*, *5-HT2B*, or *5-HT7* (Supplementary Fig. S5, C to E). These results indicate that serotonin-induced activation of the 5-HT1B, 5-HT2B, and 5-HT7 receptors in the MAG is necessary for post-mating SP replenishment.

We next used the cell type-specific drivers *Ovu-GAL4* (Chapman et al., 2003) and *D1-GAL4* (Gligorov et al., 2013) to determine whether these serotonin receptors function in main cells or secondary cells, respectively. Main cell-specific knockdown of *5-HT7* caused a modest but significant reduction in *SP* mRNA in experienced males (Fig. 3H). In contrast, because main cells normally express neither *5-HT1B* nor *5-HT2B*, their main cell-specific knockdown did not affect *SP* mRNA induction (Fig. 3H). In secondary cells, *D1-GAL4*-induced knockdown of *5-HT1B*, *5-HT2B*, or *5-HT7* each suppressed the post-mating increase in *SP* mRNA (Fig. 3I). These data suggest that serotonin acts on both main and secondary cells to regulate SP replenishment.

### Serotonin receptor-dependent signaling in MAG main and secondary cells

Next, we employed live imaging techniques to further understand the role of serotonin and serotonin receptors in main and secondary cells. 5-HT1B, 5-HT2B, and 5-HT7 are all G-protein-coupled receptors. Upon stimulation, 5-HT1B and 5-HT7 decrease and increase intracellular cAMP levels, respectively. 5-HT2B increases intracellular Ca^2+^ levels (Rosikon et al., 2023) (Supplementary Fig. S6A). Thus, we used *GAL4* lines active in the main cells or secondary cells to drive expression of the genetically encoded intracellular Ca^2+^ indicator *GCaMP6s* (Chen et al., 2013) and the intracellular cAMP indicator *Pink Flamindo* (Kurogi et al., 2023). We cultured dissected MAGs with or without serotonin and then recorded fluorescence intensities using time-lapse live imaging (Supplementary Fig. S6B). To confirm the specificity of the fluorescent reporters, we added the adenylyl cyclase activator NKH477 and the calcium ionophore ionomycin to the MAG culture. As expected, NKH477 increased Pink Flamindo fluorescence and ionomycin elevated GCaMP6s fluorescence in both main and secondary cells (Supplementary Fig. S7, A–D). Thus, we validated the functionality of both indicators in each cell type.

Under our experimental conditions, serotonin administration significantly increased cAMP levels in main cells and Ca^2+^ levels in secondary cells (Supplementary Fig. S6, C–C’’ and D–D’’), while Ca^2+^ levels in main cells remained unchanged (Supplementary Fig. S6, E– E’’). More unexpectedly, despite expression of both *5-HT1B* and *5-HT7*, we did not observe any changes in cAMP levels in secondary cells (Supplementary Fig. S6, F–F’’).

We next asked whether the intracellular cAMP and Ca^2+^ responses we observed required 5-HT7 in main cells and 5-HT2B in secondary cells, respectively. We did not test 5-HT1B in this study because serotonin administration did not change cAMP levels in secondary cells. When we knocked down *5-HT7* expression specifically in main cells, we observed a near-complete loss of serotonin-induced cAMP elevation (Supplementary Fig. S6, G–G’’). In addition, when we knocked down *5-HT2B* in secondary cells, we observed a partial but significant suppression of the serotonin-induced Ca^2+^ increase (Supplementary Fig. S6, H–H’’). Together, these results demonstrate that serotonin activates distinct second messenger pathways in main and secondary cells, consistent with their involvement in the post-mating upregulation of *SP* transcription.

### SP and SPR are required for SP replenishment in males

Our findings raised the question of how mating activates serotonin signaling to trigger SP replenishment. In general, humoral factors, particularly hormones, regulate their own biosynthesis. We hypothesized that SP itself and its receptor SPR may be involved in promoting SP replenishment, constituting a potential self-reinforcing regulatory pathway. Indeed, we found significant suppression of SP replenishment in *SPR^attP^* mutants at the protein (Fig. 4A; compare lane 4 with lane 8; also see Fig. 4, B and C) and mRNA levels (Fig. 4, D–F). We also observed this suppression of the mating-induced increase in *SP* mRNA levels in a second independent loss-of-function *SPR* mutant *SPR^Df^* (Fig. 4G). Moreover, we examined post-mating *SP* mRNA levels in the *SP^0^* line, which carries point mutations introducing a premature stop codon in the signal peptide-coding region of *SP* (Liu and Kubli, 2003) but still transcribes the entire *SP* mRNA, permitting mRNA quantification. We observed a suppression of the expected increase in *SP* mRNA levels in mated *SP^0^* males (Fig. 4H). Nevertheless, global MAG protein synthesis increased after mating in *SP^0^* males (Supplementary Fig. S8). Thus, SP is not required for the global mating-induced increase in MAG protein synthesis, despite its requirement for the normal post-mating increase in *SP* mRNA.

**Figure 4.**
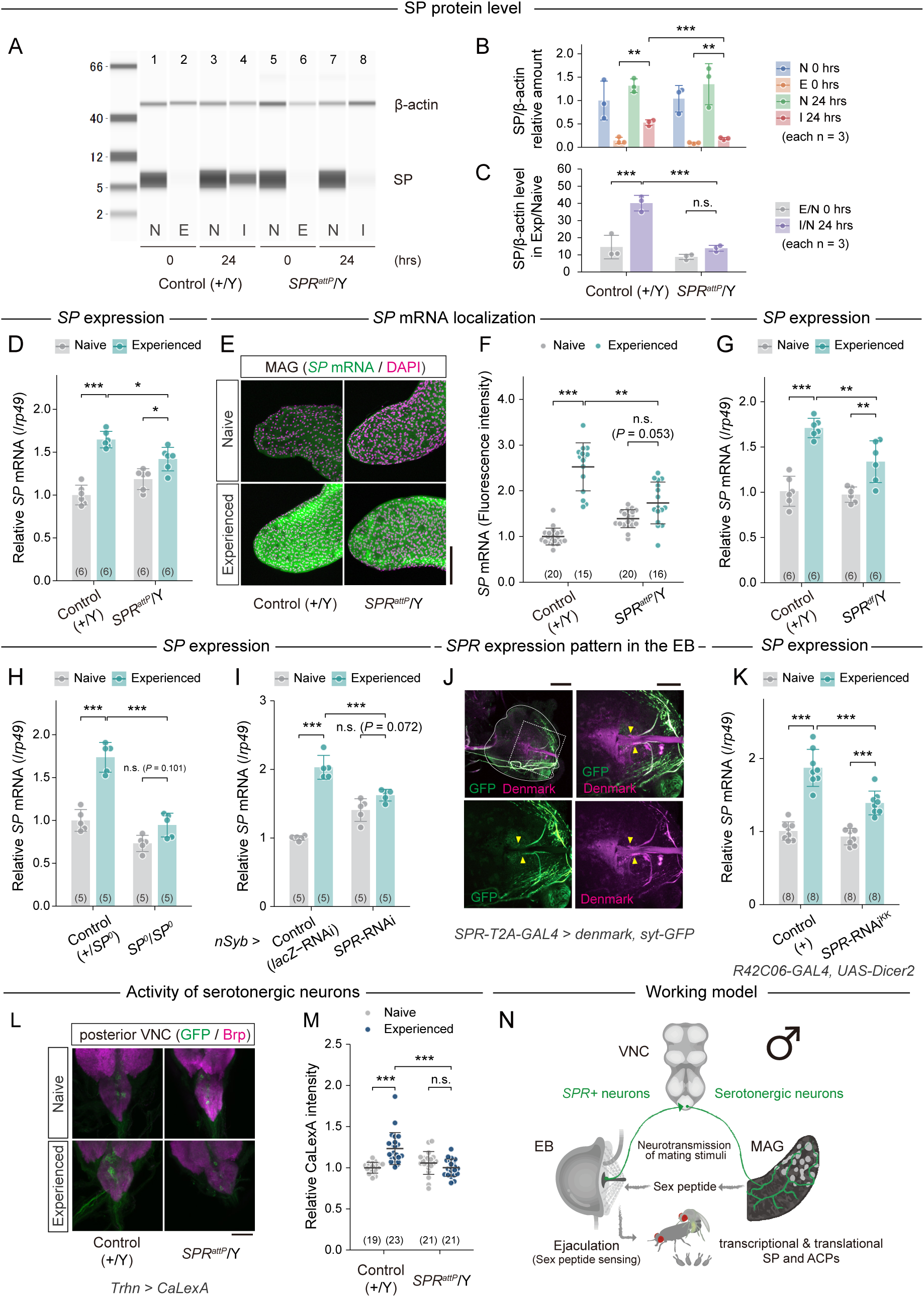
*SPR^+^*EB neurons are candidates for inducing post-mating *SP* replenishment. (A–C) Temporal changes in SP protein levels in the *SPR* mutant (*SPR^attP^*/Y) and control (*+*/Y) male reproductive tracts after mating. (A) Representative western blot image using anti-SP (7 kDa) and anti-β-actin (47 kDa) on reproductive tract samples from naive and experienced males of the indicated genotypes. Lane numbers are indicated above each lane. (B and C) Quantification of male reproductive tract SP protein levels normalized to β-actin. The relative amount of SP was normalized to the mean value of the naive control males at 0 hr. (D) *SP* mRNA levels in the male reproductive tracts of *SPR* mutant (*SPR^attP^*/Y) and control (*+*/Y) males before and after mating with values measured via RT-qPCR. *SP* mRNA levels were calculated using *rp49* as the reference gene and subsequently normalized to the mean level in naive control males. (E and F) *SP* mRNA levels before and after mating in the MAG of *SPR* mutant (*SPR^attP^*/*Y*) and control (*+*/Y) males visualized via *in situ* HCR. (E) Representative images of *SP* mRNA (green) and DAPI (magenta) staining in the tip of the MAG. Scale bar, 100 μm. (F) Quantification of *SP* mRNA intensity levels in the MAG. Signal intensities were normalized to the mean signal intensity of control males. (G–I) *SP* mRNA levels before and after mating for male reproductive tracts of the indicated genotypes. Values were measured by RT-qPCR. *SP* mRNA levels were calculated using *rp49* as the reference gene and subsequently normalized to the mean level in naive control males. (G) *SPR* loss-of-function mutant (*SPR^df^*/Y) and its control (*+*/Y). (H) *SP* knockout mutant (*SP^0^/SP^0^*) and its control (*+/SP^0^*). (I) *nSyb-GAL4* (pan-neuronal) *SPR* knockdown (*nSyb > SPR-RNAi^KK^*) and a control (*nSyb > LacZ-RNAi*). (J) *SPR-T2A-GAL4* driving GFP labeling of axons (syt-GFP) and RFP labeling of dendrites (DenMark) in the EB. The EB is outlined with the solid line in the upper left photo. The other three photos are magnified views of the area within the dotted line. Yellow arrowheads indicate cell bodies. Scale bar, whole EB: 100 μm, magnified views: 50 μm. (K) *SP* mRNA levels before and after mating in reproductive tracts isolated from males with *R42C06-GAL4* driving *SPR* knockdown (*R42C06 > SPR-RNAi^KK^, UAS-Dicer2*) and control males (*R42C06 > UAS-Dicer2*). *SP* mRNA levels were measured by RT-qPCR and normalized to *rp49*, and then normalized to the mean level in naive control males. (L and M) Pre- and post-mating CaLexA signal in serotonergic neurons of the posterior VNC in *SPR* mutants (*SPR^attP^*) and their controls. (L) Representative images of anti-GFP (green) and anti-Brp (magenta) immunostaining. Scale bar, 30 μm. (M) Quantification of CaLexA signals. Signal intensities were normalized to the mean signal intensity of control males. (N) Working model: Male mating stimuli are transmitted to the MAG via SP–SPR–serotonin signaling. All quantitative data are presented as means ± SD. The number of samples (n) is indicated in each graph. Statistical analysis: unpaired two-tailed Student’s *t*-test for (B); Tukey-Kramer’s HSD test for (C, D, G, H, I, and K); Steel-Dwass’s multiple comparison test for (F and M). \**P* < 0.05, \*\**P* < 0.01, \*\*\**P* < 0.001; n.s., not significant (*P* ≧ 0.05).

SP is not the only ligand for SPR; SPR also binds the seminal fluid protein Ductus ejaculatorius peptide 99B (Dup99B) (Saudan et al., 2002) and the neuropeptide Myoinhibiting peptide (Mip) (Kim et al., 2010). We concluded under our experimental conditions, however, that these ligands are not essential for the post-mating increase in *SP* mRNA because we observed increases in *SP* mRNA levels in both *Dup99B* and *Mip* loss-of-function mutants (Supplementary Fig. S9, A–D). Additionally, although Methuselah (Mth) was proposed as a potential SP receptor (Ja et al., 2009), *mth* loss-of-function males showed normal levels of post-mating SP replenishment (Supplementary Fig. S9E). These results support a specific requirement for SP and SPR in SP replenishment.

### Male SPR-expressing neurons similar to female SP sensory neurons may regulate post-mating SP replenishment

*SPR* is expressed in the male nervous system (Oh et al., 2014; Yapici et al., 2008). Pan-neuronal *SPR* knockdown with the *nSyb-GAL4* driver caused a significant suppression of the post-mating increase in *SP* mRNA (Fig. 4I), supporting the idea that neuronal SPR is also involved in post-mating SP replenishment.

Among *SPR*-expressing neurons in the male nervous system, we suspected a particular population of neurons comprising 2–4 cell bodies in the ejaculatory bulb (EB). We confirmed that *SPR-T2A-GAL4*, a *GAL4* transgene fused near the C-terminal region of *SPR* (Deng et al., 2019), could drive expression of *GFP* in these neurons (Fig. 4J). We will thus refer to these neurons as *SPR*^+^ EB neurons. We focused on the *SPR*^+^ EB neurons in males because they share three characteristics with f-SPSNs.

First, like f-SPSNs whose cell bodies are located near the uterus and project to the VNC (Häsemeyer et al., 2009), *SPR*^+^ EB neurons have cell bodies associated with the male reproductive tract (Fig. 4J). *R40A05-GAL4*-driven labeling revealed EB-associated neuronal cell bodies and processes extending between the EB and VNC (Supplementary Fig. S10A). This pattern is similar to that of other previously described central nervous system-projecting EB neurons (Bairati, 1968). Second, like f-SPSNs, *SPR*^+^ EB neurons are cholinergic and labeled by the cholinergic neuron-specific driver *ChAT-GAL4* (Supplementary Fig. S10B). Third, like f-SPSNs (Rezával et al., 2012), *SPR*^+^ EB neurons express the essential sex-determination genes *doublesex* (*dsx*) and *fruitless* (*fru*). Using a FLP/FRT strategy that combined *fru-FLP* and *dsx-FLP* (Rezával et al., 2014; Yu et al., 2010) with *SPR-T2A-GAL4,* we confirmed this third point with evidence that *SPR^+^* EB neurons co-express *fru* and *dsx* (Supplementary Fig. S10, C and D). Because ejaculate passes through the EB, we examined SP localization using an *SP* promoter-driven SP::GFP fusion protein (Villella et al., 2006). After mating, SP::GFP decreased in the MAG and was observed in the ejaculatory duct and EB, where it persisted for up to 24 hrs in a substantial proportion of males (Supplementary Fig. S11, A–D).

To investigate the functional role of *SPR*^+^ EB neurons, we used *R42C06-GAL4*, which labels a subset of EB neurons as well as small populations of neurons in the VNC, to knock down *SPR* (Supplementary Fig. S11, E and F). This manipulation suppressed the post-mating increase in *SP* mRNA levels (Fig. 4K), indicating that SPR in *R42C06-GAL4*-positive neurons is required for SP replenishment. In contrast, expression of membrane-tethered SP using *R42C06-GAL4* did not increase *SP* mRNA (Supplementary Fig. S11G), suggesting that providing SP to these neurons alone is insufficient. Additional mating-dependent signals or processes may therefore cooperate with the SP–SPR interaction.

We then asked whether SPR is required for mating-induced activation of MAG-projecting serotonergic neurons (Masuyama et al., 2012). In males with a wild-type background, we found significantly higher levels of CaLexA-dependent GFP fluorescence in the posterior VNC serotonergic neurons of experienced males than in naive males (Fig. 4, L and M; Supplementary Fig. S11H). In contrast, the posterior VNC serotonergic neurons of *SPR^attP^* males did not show the same increase in activity post-mating as those of control males (Fig. 4, L and M; Supplementary Fig. S11H), suggesting that SPR is essential for the mating-induced activation of MAG-projecting serotonergic neurons. Thus, we propose that mating-induced SP replenishment in the MAG is regulated by inter-tissue neural communication between the EB, VNC, and MAG (Fig. 4N).

### SP–SPR–serotonin signaling regulates the post-mating upregulation of *CG43145*

We next asked whether SP–SPR signaling regulates other candidate serotonin-dependent ACP genes identified by RNA-seq. Among the 18 ACP genes significantly upregulated after mating in controls but not in *Trhn*-null mutants, we selected seven for further analysis: *Accessory gland protein 63F* (*Acp63F*), *CG12828*, *CG31515*, *CG34130*, *CG43145*, *Male-specific transcript 57Dc* (*Mst57Dc*), and *Odorant-binding protein 56f* (*Obp56f*). Among these seven genes, *Acp63F*, *CG43145*, *Mst57Dc*, and *Obp56f* showed significantly lower expression in experienced *SP*-null males than in experienced control males by qPCR analysis (Supplementary Fig. S12A).

In contrast, the same seven genes did not show a consistent decrease in *Dup99B*-null males compared with controls (Supplementary Fig. S12B). Of these four genes, only *CG43145* was significantly reduced when *SPR* was knocked down using *R42C06-GAL4* (Supplementary Fig. S13). These results suggest that SP–SPR–serotonin signaling regulates the post-mating upregulation of *CG43145* in addition to SP.

### SP does not appear to act directly on the MAG

After detecting *SPR-T2A-GAL4*-driven fluorescence in a subpopulation of MAG secondary cells (Supplementary Fig. S14A), we wondered whether these cells may detect SP to regulate SP replenishment. When we knocked down *SPR* in the MAG using the *Prd.F-GAL4* driver (Xue and Noll, 2002), however, we still observed the post-mating increase in *SP* mRNA levels (Supplementary Fig. S14B). Moreover, using the *ex vivo* culture system, we found that addition of synthetic SP alone did not alter *SP* mRNA levels in male reproductive tracts from either control or *SP^0^*males (Supplementary Fig. S14C). Notably, we did still observe serotonin-induced upregulation of *SP* mRNA *ex vivo* in the MAG of *SPR^attP^* mutants (Supplementary Fig. S14D). These results suggest that although *SPR* is likely expressed in the MAG, SP does not directly act on the MAG to regulate post-mating SP replenishment.

### SP replenishment in males occurs widely in Sophophora species

Finally, we asked about the evolutionary conservation of post-mating SP replenishment. We focused on 12 drosophilid species, including *Scaptodrosophila lebanonensis*, a species belonging to an early-diverging lineage that retains the *SP* gene (Hopkins et al., 2024), as well as 11 additional species in which the presence or absence of female responsiveness to SP has been examined (Tsuda et al., 2015). Of the 11 species examined, nine belong to the subgenus Sophophora, including *D. melanogaster*, *D. simulans*, *D. sechellia*, *D. yakuba*, *D. erecta*, *D. ananassae*, *D. pseudoobscura*, *D. persimilis*, and *D. willistoni*. The remaining two species, *D. mojavensis* and *D. virilis*, belong to the *virilis*–*repleta* radiation. Post-mating *SP* mRNA elevation was prominent in Sophophora species, whereas it was weak or absent in species of the *virilis*–*repleta* radiation and in *S. lebanonensis* (Fig. 5 and Supplementary Fig. S15A). We also generated transgenic *D. melanogaster* lines carrying *SP* promoter sequences from *D. pseudoobscura*, *D. willistoni*, and *S. lebanonensis* fused to the GFP coding sequence (Supplementary Fig. S15B). We observed post-mating increases in *GFP* expression in each of these lines (Supplementary Fig. S15, C–E), suggesting that the heterologous *SP* promoters retain responsiveness to mating-associated signals in the *D. melanogaster trans*-regulatory environment. Furthermore, we found using RT-PCR that all 12 species express *SPR* in the male reproductive tract (Supplementary Fig. S15F), implying the potential for SPR-mediated SP detection in the EB. We also generated two *D. willistoni SPR* loss-of-function mutants and quantified their levels of *SP* mRNA. We observed a significant but incomplete reduction in *SP* expression in *D. willistoni* males lacking SPR (Fig. 5B and Supplementary Fig. S15G). Consistent with this observation, we also found a suppression of the mating-induced increase in the activity of the *D. willistoni SP* promoter in a *D. melanogaster SPR* loss-of-function mutant (Fig. 5C and Supplementary Fig. S15H). These results suggest that SPR plays at least a partial role in regulating SP replenishment in Sophophora species.

**Figure 5.**
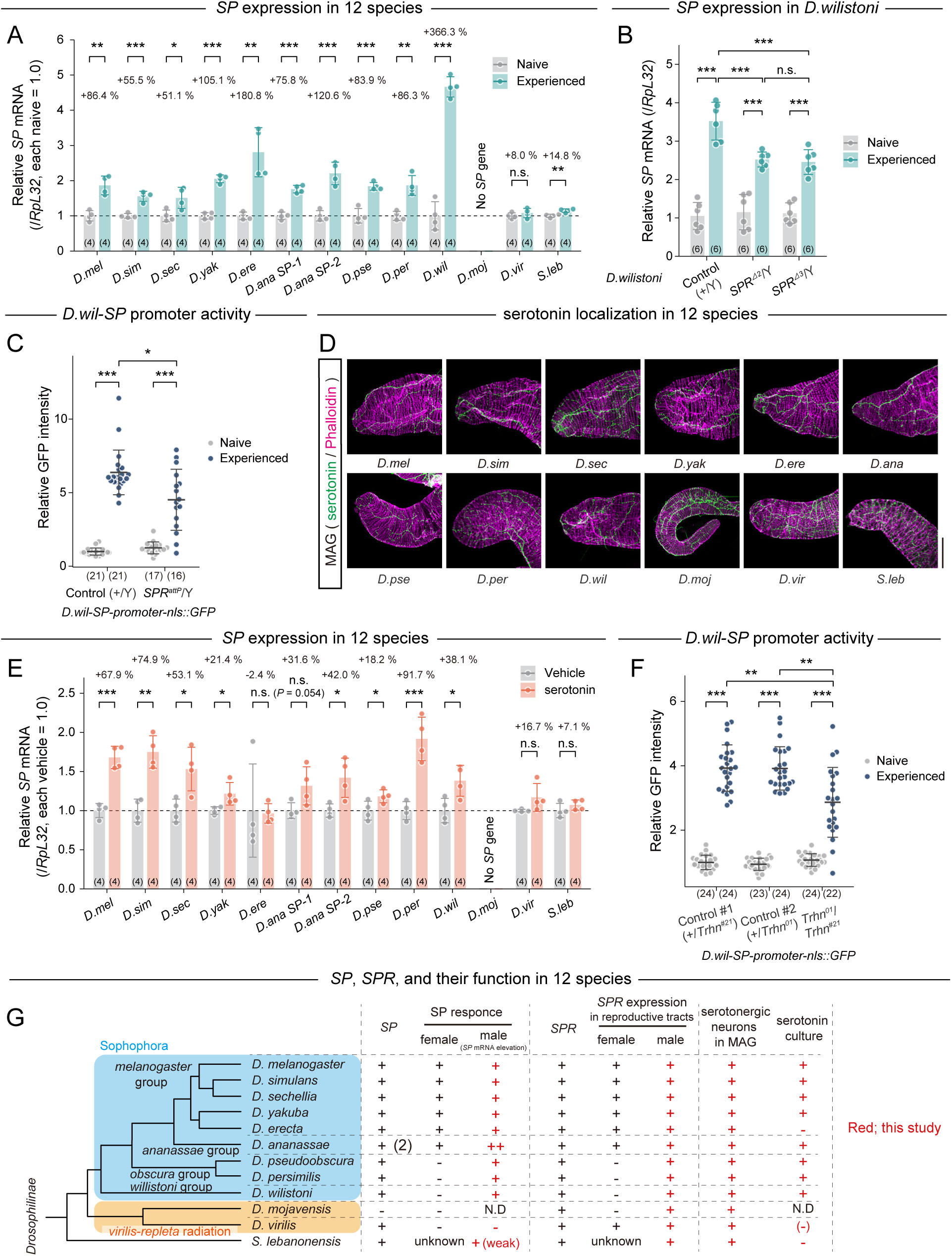
SP–SPR signaling in males extends beyond the *melanogaster* and *ananassae* groups. SP function was investigated in 12 drosophilid species: *D. melanogaster* (*D.mel*), *D. simulans* (*D.sim*), *D. sechellia* (*D.sec*), *D. yakuba* (*D.yak*), *D. erecta* (*D.ere*), *D. ananassae* (*D.ana*), *D. pseudoobscura* (*D.pse*), *D. persimilis* (*D.per*), *D. willistoni* (*D.wil*), *D. mojavensis* (*D.moj*), *D. virilis* (*D.vir*), and *S. lebanonensis* (*S.leb*). *SP* mRNA levels in male reproductive tracts isolated from each strain were measured by RT-qPCR and normalized to *RpL32* (*rp49*). (A) *SP* mRNA levels before and after mating for males of 11 of the 12 species (excluding *D.moj*). Values are shown relative to the levels of naive males from each respective species. See Supplementary Fig. S15A for a quantification of *SP* mRNA levels relative to those of naive *D. mel*. (B) *SP* mRNA levels before and after mating in the male reproductive tract of the *D.wil*-*SPR* mutants (*SPR*^Δ*2*^/Y and *SPR*^Δ*3*^/Y) and control (+). *SP* mRNA levels were normalized to the mean level in naive control males. (C) *D. wil*-*SP* promoter activity before and after mating in the *SPR* mutant and its control. The *D. wil-SP* promoter-GFP reporter transgene was introduced into *D. melanogaster*, and GFP intensity was assessed by immunostaining. Signal intensities were normalized to the mean signal intensity of control males. (D) Representative images of MAGs from the 12 species stained with anti-serotonin (green) antibody and phalloidin (magenta). Image acquisition settings for each of the 12 photos were different so that all signals would be visible. Scale bar, 100 μm. (E) *SP* mRNA levels in male reproductive tracts from 11 species after 6 hrs of *ex vivo* culture with and without serotonin. *SP* mRNA levels are shown relative to the control for each species. (F) *D.wil*-*SP* promoter activity before and after mating in the *Trhn* mutant (*Trhn^01^/Trhn^#21^*) and its controls (+/*Trhn^#21^* and +/*Trhn^01^*). The *D.wil-SP* promoter-GFP reporter transgene was introduced into *D. melanogaster*, and GFP intensity was assessed by immunostaining. Signal intensities were normalized to the mean signal intensity of control males (+/*Trhn^#21^*). (G) Schematic summarizing the function of *SP* in 12 drosophilid species. Results from Tsuda et al. (2015) are shown in black. Results from this study are shown in red. The presence or absence of an SP sequence and its function in each species is indicated by + or −, respectively. “(−)” indicates that the increase in *SP* mRNA following serotonin treatment or mating did not reach statistical significance, although the mean level in the treated or mated group was at least 10% higher than that in the corresponding control group. N.D. indicates not determined. Note that *D.moj* does not have an *SP* gene and *D. ana* has two *SP* genes. All quantitative data are presented as means ± SD. Statistical analysis: unpaired two-tailed Student’s *t*-test for (A and E); Tukey-Kramer’s HSD test for (B); and Steel-Dwass’s multiple comparison test for (C and F). \**P* < 0.05, \*\**P* < 0.01, \*\*\**P* < 0.001; n.s., not significant (*P* ≧ 0.05).

We next examined the role of serotonin in SP replenishment in these 12 drosophilid species. All 12 species had serotonin-positive neuronal projections to the MAG (Fig. 5D). To determine whether serotonin increases *SP* mRNA in the MAG in these species, we performed *ex vivo* cultures of the male reproductive tracts in the presence or absence of serotonin. Serotonin treatment increased *SP* mRNA levels in multiple Sophophora species. However, the magnitude of this effect varied considerably and may reflect lineage-specific modulation rather than a simple phylogenetic trend (Fig. 5E). In *S. lebanonensis*, serotonin treatment showed a trend toward increased *SP* mRNA that did not reach statistical significance, whereas its *SP* promoter showed a significant response in the transgenic assay (Supplementary Fig. S15, C–E). To further assess *SP*-promoter activity in Sophophora species, we used *D. melanogaster* lines carrying *D. pseudoobscura* and *D. willistoni SP* promoters fused to the coding sequence of GFP. Using *ex vivo* cultures of the resulting male reproductive tracts, we found that serotonin administration increased the activity of both the *D. pseudoobscura* and *D. willistoni SP* promoters (Supplementary Fig. S15I). Furthermore, we observed a significant suppression of the mating-induced activation of the *D. willistoni*-derived *SP* promoter in the absence of *Trhn* (Fig. 5F and Supplementary Fig. S15J). These results support the hypothesis that post-mating SP replenishment—mediated via the SP–SPR–serotonin axis—is functionally conserved in Sophophora species (Fig. 5G). Given that the typical SP-mediated PMRs of females are absent in the *obscura* and *willistoni-*group species (Tsuda et al., 2015), our results suggest that the role of SP and SPR in SP replenishment in males may be more ancestral than their role in regulating female PMRs in the evolution of Drosophilinae (see the Discussion for more details).

## Discussion

SP is a seminal fluid protein that plays a major role in inducing female PMRs. Although males do display some PMRs, such as the replenishment of ACPs in the MAG, the mechanisms that trigger these responses in males remain poorly understood. In this study, we show that SP and SPR play a previously unknown role in the regulation of post-mating SP replenishment in males. This process involves inter-tissue communication between the EB, VNC, and MAG through SP–SPR–serotonin signaling. One of our key findings is that SP replenishment in males requires SP–SPR signaling. Although previous studies identified SPR-independent functions of SP in males, including courtship behavior and lipid microcarrier formation (Denis et al., 2018; Wainwright et al., 2021), this is the first demonstration of an SP–SPR-dependent function in males. Thus, SP not only acts on females after mating but also regulates male reproductive physiology through inter-tissue signaling.

We hypothesize that SP is received by *SPR*-expressing neurons in the ejaculatory bulb. Interestingly, Dup99B, another SPR ligand expressed in the ejaculatory duct (Rexhepaj et al., 2003), does not appear to contribute to the regulatory pathway identified in this study. Why SP and Dup99B differ in their contributions to post-mating SP replenishment remains unclear. Future studies examining their relative abundance, localization, and accessibility to SPR-expressing neurons may help clarify the basis of this difference.

Another key finding of this study is our identification of serotonin as a crucial trophic factor for enhancing the production of SP and some other ACPs in the MAG. Serotonin activates 5-HT7-dependent cAMP signaling in main cells and 5-HT2B-dependent Ca^2+^ signaling in secondary cells, consistent with their known coupling to Gαs and Gαq, respectively (Giulietti et al., 2014). While 5-HT1B couples to Gαi and suppresses intracellular cAMP levels (Giulietti et al., 2014), we did not observe this in the MAG. Although 5-HT1B would likely have an inhibitory effect on cAMP signaling, we found that *5-HT1B* RNAi in the MAG causes the same effect on SP production as *5-HT2B* or *5-HT7* RNAi. Further studies will be necessary to understand how an inhibitory signal from 5-HT1B could have a positive effect on SP replenishment. We were also interested to find that SP replenishment requires 5-HT7 in the *SP*-expressing main cells, as well as 5-HT1B, 5-HT2B, and 5-HT7 in secondary cells. Given that SP is produced primarily by the main cells (Chapman et al., 2003; Chen et al., 1988; Kubli, 2003), the serotonin signaling that occurs in the main cells seems to affect SP replenishment directly. In the secondary cells, however, serotonin signaling likely modulates SP replenishment indirectly via the main cells. This raises the possibility that main cells and secondary cells actively communicate, leading to increased SP production in main cells. From a morphological perspective, the large apical cell-cell contact zone between main cells and secondary cells may facilitate cell-cell interaction between the two cell types (Prince et al., 2019). It will be intriguing to discover the identity of the serotonin-induced signals produced and released by secondary cells, as well as how main cells receive these signals to regulate SP production after mating.

Mechanistically, we propose that *SPR^+^*EB neurons sense SP as it passes through the EB during ejaculation and relay this mating signal to MAG-projecting serotonergic neurons. Consistent with this model, SP::GFP was observed in the ejaculatory duct and EB after mating, although only in approximately half of the experienced males examined, possibly reflecting transient passage of SP through these structures. How *SPR^+^*EB neurons communicate with MAG-projecting serotonergic neurons remains unknown. Moreover, because *R42C06-GAL4* labels additional neuronal populations, we cannot exclude contributions from *SPR* expressed in other neurons. More selective genetic tools, such as appropriate *split-GAL4* drivers, will therefore be required to establish the specific role and connectivity of *SPR^+^*EB neurons. Our loss-of-function results support a requirement for SP and neuronal SPR in the normal post-mating increase in *SP* mRNA, but do not establish direct modulation of *SPR*-positive EB neurons by SP. Membrane-tethered SP did not induce this response with *R42C06-GAL4*, leaving unresolved whether additional mating-dependent signals or limitations of this manipulation account for the lack of sufficiency.

Our final key discovery is that mating- and serotonin-induced SP replenishment occurs in Sophophora species. We genetically demonstrated that SP replenishment at least partially depends on SPR function in male *D. willistoni*, a member of the subgenus Sophophora. This observation suggests that SP–SPR signaling in males is conserved in Sophophora. Notably, SP also has an SPR-independent role in microcarrier formation in males, which has been proposed as another ancestral male function of SP (Wainwright et al., 2021). Thus, SP may have had multiple ancestral functions in males. Nevertheless, our findings suggest that the SP–SPR signaling axis itself had a male role in regulating SP replenishment before its prominent role in female PMRs evolved. Consistent with this possibility, SP plays a much more limited role in modulating female PMRs in the Sophophora species outside the *melanogaster* and *ananassae* groups (Tsuda et al., 2015). We therefore propose that the well-known role of SP–SPR signaling in female PMRs represents a derived function relative to its role in males.

Male *SPR^+^* EB neurons and female f-SPSNs share several molecular and anatomical features. Both have reproductive tract-associated cell bodies and VNC projections and express *dsx* and *fru*. These similarities raise the possibility of an evolutionary relationship between the two SP-sensing circuits. However, the weak post-mating SP replenishment observed in *D. virilis* and *S. lebanonensis* leaves unresolved whether this function arose later in evolution or was secondarily reduced or lost in these lineages. Determining the functions of SP and SPR in these earlier-diverging lineages may therefore provide insight into the deeper evolutionary origins of this signaling system. It also remains unclear why Sophophora males outside the *melanogaster* and *ananassae* groups replenish SP after mating despite the apparent absence of canonical SP– SPR-mediated PMRs in females (Tsuda et al., 2015).

SP represents only one of more than 150 ACPs identified in *D. melanogaster* (McQuarrie et al., 2025), several of which are also replenished after mating (Hollis et al., 2019; Koppik et al., 2018; Sirot et al., 2009). Although we focused primarily on SP, our results indicate that SP also contributes to the post-mating upregulation of a subset of other ACP genes. Among these, however, only *CG43145* was reduced following *SPR* knockdown in *R42C06-GAL4*-positive neurons, whereas loss of *Dup99B* did not consistently affect the ACP genes examined. Thus, SP does not appear to serve as a generic mating signal controlling most or all ACPs; rather, the SP–SPR–serotonin pathway regulates SP together with a limited subset of ACPs. Identifying the downstream transcriptional mechanisms that confer this selectivity will be an important next step.

It is important to note, however, that the SP–SPR–serotonin pathway is unlikely to be the sole regulator of post-mating ACP replenishment. Indeed, SP replenishment is not completely abolished in *D. melanogaster* or *D. willistoni* lacking *SPR* function, suggesting that additional regulatory signals or pathways contribute to this response. One candidate is juvenile hormone (JH), as JH III reportedly induces protein synthesis in the *D. melanogaster* MAG *ex vivo* (Yamamoto et al., 1988). Interestingly, SP itself stimulates JH biosynthesis in the female *corpora allata ex vivo* (Moshitzky et al., 1996). Mechanical stimuli associated with changes in MAG size after mating may also contribute to the regulation of *SP* and ACP gene expression, although mechanical emptying of the *D. funebris* MAG *in vitro* does not increase seminal fluid protein production (Baumann, 1974). How these signals interact with the SP–SPR–serotonin pathway remains an important question. Although our insemination and fecundity assays do not directly quantify the amount of accessory gland secretion transferred during mating, SP levels appeared similarly depleted in experienced control, *Trhn*-null, and *SPR*-null males. This observation argues against impaired SP release as the primary explanation for the attenuated post-mating SP replenishment in these mutants, although altered transfer of other seminal fluid components cannot be excluded.

Beyond *Drosophila*, seminal fluid molecules are critical for reproduction across diverse taxa (Bromfield, 2014; Meuti and Short, 2019; Schjenken and Robertson, 2014). Although *SP* is conserved only in Drosophilinae, analogous systems for post-mating replenishment of seminal fluid proteins may exist and remain to be discovered in other iteroparous animals.

## Materials and Methods

### Wild-type *Drosophila* strains and husbandry

While Canton-S *and w^1118^* were used as wild-type in most analyses, the genome project WGS strain (k-s04, KYORIN-Fly: Drosophila species stock center) was used in the interspecies comparisons shown in Figure 5 A, D and E, and Supplementary Fig. S15A. This comparative analysis also included the following *Drosophila* species obtained from KYORIN-Fly: *D. simulans* (E-10501), *D. sechellia* (k-s10), *D. yakuba* (k-s03), *D. erecta* (k-s02), *D. ananassae* (k-s01), *D. pseudoobscura* (k-s12), *D. persimilis* (k-s11), *D. willistoni* (k-s13), *D. mojavensis* (k-s15), *D. virilis* (k-s14), and *S. lebanonensis* (E-21401).

Flies of all species were maintained on a standard diet (0.55 g of agar, 10.0 g of glucose, 9.0 g of cornmeal, 4.0 g of yeast extract, 300 μl of propionic acid, 350 μl of 10% butylparaben in 70% ethanol, and 100 ml of water). Unless otherwise stated, experiments with *D. melanogaster* were conducted at 25°C under a 12:12-hr light/dark cycle. *D. persimilis* flies were maintained at 20°C under a 16:8-hr light/dark cycle, and the rest of the *Drosophila* species were maintained at 23°C under a 16:8-hr light/dark cycle.

### Mutant and transgenic *D. melanogaster* strains

The *Drosophila* strains used in this study are listed in Table S1. We used homozygous *SP□/SP□* males, based on previous reports showing that these males can mate (Garbe et al., 2016). *R40A05-GAL4* was used to visualize EB-associated neurons and their processes (Supplementary Figs. S10A and S11E), whereas *R42C06-GAL4* was used for *SPR* knockdown (Fig. 4K and Supplementary Fig. S13). *R42C06-GAL4* also labels neuronal populations in the VNC.

Immunostaining with an anti-Crz antibody confirmed Crz proteins were undetectable in the nervous system of *Crz^10A^*/*Crz ^sk2-5^* mutant animals (Imura et al., 2020; Sha, 2013) (Supplementary Fig. S1, F and G).

### Establishing *SP-promoter GFP* reporter lines in *D. melanogaster*

Approximately 1,600–2,200 bp segments upstream of the *SP* gene from each *Drosophila* species genome were cloned into *Eco*RI-*Bam*HI-digested pS3aG (Williams et al., 2008) using the In-Fusion HD Cloning Kit (Clontech, 639648). The primers used to amplify these upstream genomic sequences are listed in Table S2. The constructs were then injected into embryos of the *phiC31* integrase strain (*nos-phiC31; attP40; attP2*). F1 males were crossed with *w^1118^; Roi/CyO; TM3 Sb/TM6 Tb Hu* females. White-positive F2 progeny carrying the *SP-promoternls::GFP* transgenes on the second chromosome were selected. Heterozygous *SP-promoternls::GFP* flies were used for observing and quantifying GFP signals.

### Establishing genetic null mutants for *Trhn*, *Henna*, and *Dup99B* in *D. melanogaster*, and for *SPR* in *D. willistoni*

The CRISPR-Cas9 system with the pBFv-U6.2 vector (Kondo and Ueda, 2013), provided by the National Institute of Genetics (NIG) in Japan, was used to generate each *D. melanogaster* mutant. One or two sgRNA target sites were selected for each gene to minimize potential off-target effects. Sense and antisense oligonucleotides corresponding to the sgRNA target sequences were annealed and cloned into *Bbs*I-digested pBFv-U6.2 vector. The resulting gRNA constructs were injected into *y¹ v¹ nos-phiC31; attP40* embryos. The progenies were then crossed to flies carrying balancer chromosomes, and mutants were identified by genomic DNA sequencing.

*D. willistoni SPR* mutants were also generated with the CRISPR-Cas9 system. Three gRNA target sites were selected in the 1^st^ exon of *SPR* (LOC6649192) (Supplementary Fig. S15G). Two different crRNAs, tracrRNA, and Cas9 protein were injected into *D. willistoni* (k-s13) embryos. Genomic PCR was used to screen F1 progeny for those with deletions in the 1^st^ exon of *SPR*. Then, the deleted regions were identified by genomic DNA sequencing (Supplementary Fig. S15G). The *SPR*^Δ*2*^ and *SPR*^Δ*3*^ mutants were maintained as homozygotes.

The sequences of the gRNAs or oligonucleotides used for cloning the gRNAs are listed in Supplementary Table S3.

### Transgenic manipulation of neuronal activity

We manipulated neuronal activity by pairing *Kir2.1*, *shibire^ts^*(*shi^ts^*), or *TrpA1* with *GAL4* drivers labeling neurons of interest. Flies expressing *Kir2.1*, which constitutively suppresses neuronal excitability via potassium channel overexpression (Baines et al., 2001), were maintained at 25°C. Flies expressing *shi^ts^*, which blocks synaptic transmission at the restrictive temperature (Kitamoto, 2002), were shifted to 30°C one day prior to mating. In contrast, flies expressing *TrpA1*, which allows temperature-dependent neuronal activation (Alekseyenko et al., 2010; Hamada et al., 2008), were exposed to 30°C for one hr immediately before mating to induce neuronal activity. Naive males with *shi^ts^* and *TrpA1* were collected after eclosion and then reared at 20°C for 1 week before experiments.

### Mating assays

Adult male flies were reared at 25°C for 5–7 days and then transferred to new vials at a male-to-female ratio of 1:3, where they were allowed to mate freely for 24 hrs. Because males transfer approximately one-third of the MAG contents during a single copulation (Herndon et al., 1997), this mating regime is expected to result in substantial depletion of accessory gland contents.

This study also used homozygous *SP□/SP□* males. Under our experimental conditions, these males were capable of successful mating and were therefore used directly for all mating-based assays.

### Immunohistochemistry and microscopy

Brains, ventral nerve cords (VNCs), and reproductive tracts were dissected in Schneider’s *Drosophila* medium (Thermo Fisher Scientific, 21720024) and fixed in 4% formaldehyde in PBS for 30 to 60 min at room temperature. Fixed samples were washed three times in PBS. Samples were then blocked in a blocking solution (PBS with 0.1% Triton X-100 and 2% bovine serum albumin) for 1 hr at room temperature and incubated with a primary antibody in blocking solution at 4°C overnight. The primary antibodies used in this study were rabbit anti-Crz (a gift from Jan A. Veenstra; 1:1000) (Veenstra, 1991), rabbit anti-serotonin (Sigma-Aldrich, S5545; 1:1000), mouse anti-Fas3 (Developmental Studies Hybridoma Bank (DSHB), 7G10; 1:50), mouse anti-Chaoptin (Chp) (DSHB, 24B10; 1:50) (Kroeger, 2017), mouse anti-Bruchpilot (Brp) (DSHB, nc82; 1:50), rabbit anti-RFP (Abcam, ab62341; 1:1000), and chicken anti-green fluorescent protein (GFP; Abcam ab13970; 1:8000). After washing, the samples were incubated with fluorophore-conjugated secondary antibodies (Alexa Fluor 488, 546, or 555; Thermo Fisher Scientific) and Alexa568-conjugated phalloidin (Thermo Fisher Scientific, A12380) diluted 1:200 in blocking solution for 2–3 hrs at room temperature or for at least 16 hrs at 4°C. The tissues were then washed again before nuclear staining with 4′,6-diamidino-2-phenylindole (DAPI; Thermo Fisher Scientific, 62247; 1:10,000) for 15 min. After another washing step, all samples were mounted in FluorSave reagent (Merck Millipore, 345789).

Samples were visualized using a Zeiss LSM 700 confocal microscope, Zeiss LSM 900 confocal microscope, Zeiss AxioPlan 2, or Leica Thunder Imaging System. Fluorescence intensity in confocal sections was measured using ImageJ.

### Quantification of fluorescence intensity

Fluorescence intensity was quantified using FIJI/ImageJ (https://imagej.net/). For experiments with MAGs, regions of interest (ROIs) were defined in both lobes, and the mean fluorescence intensity was used as the representative value for each MAG for subsequent plotting and statistical analysis.

To quantify *SP* mRNA levels from isHCR images, ROIs in the distal tips of the glands were manually selected, and the mean fluorescence intensity in each ROI was measured. The same quantification procedure was applied to OPP staining images to assess nascent protein synthesis.

For *SP*-promoter-GFP reporter analysis, GFP fluorescence intensity was quantified in the distal tips of the glands by identifying nls::GFP-positive nuclei. Mean GFP intensity was measured across multiple nuclei using the “Analyze Particles” function of ImageJ, and the resulting values were normalized to those obtained under control conditions.

Serotonin immunoreactivity was quantified in the distal tips of MAGs by defining ROIs corresponding to serotonin-positive regions. Total signal intensity was calculated as the product of mean fluorescence intensity and ROI area, and this value was normalized to the area of each MAG distal tip region.

For the CaLexA experiments in the VNC, reporter activity was quantified by defining ROIs corresponding to the GFP signal of each animal. Mean fluorescence intensity was measured for each ROI and normalized to control conditions for statistical comparisons.

### *In situ* hybridization chain reaction in the male reproductive tract

*In situ* hybridization chain reaction was performed as previously described (Mikami et al., 2025) with minor modifications. Male reproductive tracts were dissected in Schneider’s *Drosophila* medium and fixed in 4% formaldehyde in PBS for 30 min at room temperature. Fixed samples were washed three times in PBS with 0.1% Triton X-100. The fixed samples were rinsed sequentially in mixtures of methanol and PBS containing 0.1% Triton X-100 at methanol-to-buffer ratios of 1:3, 1:1, and 3:1, and then stored in 100% methanol at −20°C for periods ranging from 1 day to 1 week. After that, samples were sequentially rinsed in 3:1, 1:1, and 1:3 mixtures of methanol and SSCT (0.1% Tween-20 in 1× SSC), and then finally washed three times in SSCT. Pre-hybridization was conducted in hybridization buffer (15% ethylene carbonate, 5× SSC, 0.1% Tween-20, and 50 μg/ml heparin in 1× Denhardt’s solution) at 45°C for 30 min. Hybridization was performed at 45°C overnight using probe sets diluted in hybridization buffer (two sets of SP probes at 20 nM). The probes were designed semi-automatically using a published R pipeline (Kozono et al., 2026) (See Table S4). Samples were then washed three times with SSCT, followed by a rinse in a 1:1 mixture of hybridization buffer and SSCT. After these post-hybridization washes, hairpin DNA pairs (SaraFluor™488-S45) were annealed and applied to the samples in amplification buffer (Nepagene, IPL-AB) for 2 hrs at 25°C. Finally, the samples were washed in SSCT containing DAPI and mounted in FluorSave reagent. The samples were stored at low temperature and photographed with an LSM700.

### Quantitative and non-quantitative reverse transcription PCR

To quantify *SP* mRNA levels in the various *Drosophila* species, reproductive tracts of 8–10 adult male flies (5–8 days after eclosion) were dissected for each sample. Total RNA was extracted using the RNAiso Plus reagent (TaKaRa, 9109). Then, cDNA was prepared with ReverTra Ace qPCR RT Master Mix with gDNA Remover (TOYOBO, FSQ-301). RT-qPCR was performed using THUNDERBIRD NEXT SYBR qPCR Mix (TOYOBO, QPX-201) with a Thermal Cycler Dice TP800 system (TaKaRa). In this study, we performed both absolute and relative (^ΔΔ^Ct method) quantification, depending on the plasmid preparation used for the standard curve. For absolute quantification, serial dilutions of plasmids containing the open reading frame of each gene and occasionally the untranslated region were used as standards. The amount of target mRNA was normalized to *ribosomal protein 49* (*rp49*)/*ribosomal protein L32* (*RpL32*) and relative fold changes were calculated. The primers used in this study are described in Tables S5 and S6. The primers for *rp49* and *SP* in *D. melanogaster* were previously described (Chapman et al., 2003; Tsuda et al., 2015).

Non-quantitative RT-PCR was conducted to determine whether the *SPR* gene is expressed in various *Drosophila* species. Tissues including the MAG, ejaculatory duct, and EB of 10 adult male flies (5–7 days post-eclosion) were dissected for each sample. Total RNA was extracted using the RNAiso Plus reagent, and cDNA was prepared using ReverTra Ace qPCR RT Master Mix with gDNA Remover. RT-PCR was performed using KOD-Plus-Neo (TOYOBO, KOD-401) with a T100 Thermal Cycler system (Bio-Rad). Amplicons were subjected to electrophoresis on a 2% agarose gel to confirm bands of the target length. Amplicons were subcloned into pBluescript II (SK-) plasmid, followed by sequencing to confirm the amplicons corresponded to *SPR* in each *Drosophila* species (except for *S. lebanonensis*). The primers used for these experiments are described in Table S6.

### RNA-seq

The RNA-seq transcriptional data of adult reproductive glands obtained from each genotype used for Supplementary Fig. S3, A and B, are available from DNA Data Bank of Japan Sequence Read Archive (BioProject No. PRJDB40147). RNA purification, RNA quality check, library synthesis, library quality check, RNA sequencing, and RNA data analysis were conducted by Tsukuba i-Laboratory LLP (https://www.tsukuba-i-lab.com/), according to their protocol and workflow (https://www.md.tsukuba.ac.jp/basic-med/genome/iLab/RNAseq_workflow_v1.pdf). In this study, an average of 42 million reads per biological replicate was obtained. FastQC was used to evaluate the quality of the raw, paired-end reads. Among these data, 155 representative ACPs, as defined by a previous study (McQuarrie et al., 2025), were selected.

To evaluate which ACP genes were differentially expressed, differential expression analyses were conducted independently for two comparisons: (1) naive control males versus experienced control males; (2) naive *Trhn*-null mutant males versus experienced *Trhn*-null mutant males. *P* values and fold change (FC) values were calculated using the Exact test for differential gene expression (EDGE) test implemented in CLC software. For comparison of gene expression among 155 ACP genes, *P* values were corrected using the Benjamini–Hochberg false discovery rate (FDR). Fold changes calculated using CLC were represented as simple ratios when gene expression was increased. When gene expression was decreased, the inverse of the ratio was taken and assigned a negative sign. These values were provided in the Source Data file for each RNA-seq experiment.

For heatmap visualization, normalized values were log_2_-transformed after adding 1 to avoid zero values. The normalized data from the four experimental groups were used for analysis. Heatmaps were generated using hierarchical clustering in Morpheus (Broad Institute; https://software.broadinstitute.org/morpheus/) with one minus Pearson correlation as the distance metric.

### Adult fly injection

A protocol to inject solutions into naive males was described previously (Ameku et al., 2018). We utilized 1 mM serotonin (Nacalai tesque, 18961-41) for injection as previously described (Tayler et al., 2012). Then, reproductive tracts were dissected 3–16 hrs after serotonin injection.

### Preparation of 5-HTP containing diet

The pharmacological protocol used in this study was previously described (Dierick and Greenspan, 2007; Hu et al., 2020) with some modifications. Newly emerged naive males were collected and reared on standard food for 5 days. They were then reared for 3 days on a standard diet mixed with 5-HTP (Sigma-Aldrich, H9772) at a final concentration of 50 mM. Thereafter, males were mated with females for 24 hrs under standard feeding conditions with the addition of 5-HTP.

### Western blot assay and sample preparation

The Simple Western system from Abby (Bio-Techne) was used for capillary western blotting. To examine SP protein levels, the reproductive tracts of 10 adult male flies (5 days after eclosion) were dissected for each sample. The samples were homogenized in 40 μL 1× SDS sample buffer with 1/20th the volume of 2-mercaptoethanol and then incubated at 98°C for 5 min. The levels of SP and β-actin proteins were then quantified using Abby with 1/10 diluted samples. For SP quantification, rabbit anti-SP antibody (a gift from Mariana F. Wolfner; 1:200-400) (Misra and Wolfner, 2020), the EZ Standard Pack 5 (Bio-Techne, PS-ST05EZ-8), and a 2– 40 kDa Separation Capillary Cartridge (Bio-Techne, SM-W012-01) were used. For β-actin quantification, mouse anti-β-actin (Santa Cruz Biotechnology, sc-47778, 1:50), the EZ Standard Pack 1 (Bio-Techne, PS-ST01EZ-8), and a 12–230 kDa Separation Capillary Cartridge (Bio-Techne, SM-W004-01) were used. SP and β-actin were detected with the anti-rabbit (Bio-Techne, DM-001) and anti-mouse detection modules (Bio-Techne, DM-002), respectively.

### Egg-laying assay

Flies were reared at 25 °C and aged for 5 days. Three virgin females were allowed to mate with a single target-genotype male for 24 hrs in a food-containing 35-mm Petri dish (for “Day 1” mating in Fig. 2H). After 24 hrs, the females were transferred to a fresh food-containing Petri dish, and the male was transferred to another dish containing standard food together with three fresh virgin females (for “Day 2” mating in Fig. 2H). After an additional 24 hrs, the females used for the second round of mating were also transferred to a fresh food-containing Petri dish. For both sets of females, egg laying was quantified over 48 hrs—24 hrs in the presence of the male followed by 24 hrs of isolation.

### SP refolding and purification

The SP refolding procedure, which involved formation of an intramolecular disulfide bond within an SP peptide, was performed as previously described (Tsuda et al., 2015; Yapici et al., 2008). In brief, *D. melanogaster* SP (NH□-WEWPWNRKPTKFPIPSPNPRDKWCRLNLGPAWGGRC-COOH) was synthesized artificially by Eurofins Genomics, Inc. The synthetic peptide was initially dissolved in a solution containing 3% (v/v) dimethyl sulfoxide (DMSO) and 10 mM ammonium bicarbonate. The solution was incubated for more than 36 hrs to allow refolding. The solution was then lyophilized to completely remove the solvents. The dried peptide sample was re-dissolved in Milli-Q water, then centrifuged at 18,300 *x g* for 10 min to remove insoluble materials. The resulting supernatant was then mixed with an equal volume of a solution containing 0.1% trifluoroacetic acid (TFA) and 5% acetonitrile (AcCN), and subjected to reversed-phase high-pressure liquid chromatography (RP-HPLC) using a CAPCELL PAK C18 column (SG120, 3 mm × 150 mm, SHISEIDO, Japan). The peptide was eluted using a linear gradient of AcCN from 5% to 75% over 65 min in 0.1% TFA, at a flow rate of 0.5 ml per min. During fractionation, the presence of the peptide in the eluent was monitored at 190 nm. After lyophilization of the peptides, the aliquots of each fraction were subjected to non-reducing SDS-PAGE to check whether the fractions contained monomeric or multimeric forms of SP. The fractions with only the monomeric SP were stored at -80 °C until use.

### *Ex vivo* male reproductive tract culture

*Ex vivo* reproductive tract culture experiments were performed as described (Ameku et al., 2018; Yoshinari et al., 2020). Adult male reproductive tracts were dissected and collected in Schneider’s *Drosophila* medium. Ten male reproductive tracts were transferred to 35-mm plastic dishes containing 3 ml of Schneider’s Drosophila medium supplemented with 10% fetal bovine serum and 1× Penicillin–Streptomycin Solution (Fujifilm-Wako, 168-23191). The culture medium was supplemented with either serotonin (Nacalai Tesque, 18961-41; final concentration, 1 mM), synthetic SP (final concentration, 1 μM), or control vehicle (Schneider’s Drosophila medium for serotonin and distilled water for SP). The cultures were then incubated at 25°C for 3–16 hrs, after which the samples were homogenized for *SP* mRNA quantification.

### *Ex vivo* MAG live imaging assay

Adult naive males were maintained on standard diet until 4–8 days post-eclosion and then dissected in Schneider’s *Drosophila* medium. Adult knockdown flies for the *5-HT2B*-RNAi experiment (Supplementary Fig. S6H) were reared at 23°C because no adults emerged at 25°C. The dissected tissues were kept in a glass-bottom dish (35×10 mm, IWAKI, 3910-035) with an insect pin ( 0.10 mm, Ento Sphinx Insect Pins) and silicone grease (Beckman, 335148). Then, 20 μL of Schneider’s Drosophila medium was added to cover the tissue. Live imaging was performed at 25°C using a laser scanning confocal microscope (LSM900, Carl Zeiss) with a 40× objective lens (water-immersion). Time-lapse images were acquired every 2 sec. The imaging began without any supplements. After the 49^th^ or 50^th^ frame was captured (i.e., 96 or 98 sec after imaging began), the time-lapse recording was paused for the addition of 80 μL of SDM with or without 1.25 mM Serotonin (Nacalai tesque, 18961-41; final concentration: 1 mM), 250 nM ionomycin (Abcam, AB120370; final concentration: 200 nM), or 125 μM NKH477 (Sigma-Aldrich, N3290; final concentration: 100 μM). Then, the time-lapse recording was resumed, and the time of supplement addition was defined as t = 0. To examine Ca levels, GCaMP6s and mCD8::RFP were excited with 488 nm and 561 nm lasers, respectively. To measure intracellular Ca^2+^ levels, GCaMP6s and mCD8::RFP were excited using 488 nm and 561 nm lasers, respectively. For cAMP measurements, mCD8::GFP and Pink Flamindo were excited using 488 nm and 561 nm lasers, respectively. Fluorescence intensity ratios were calculated as GCaMP6s/mCD8::RFP for Ca^2+^ measurements and Pink Flamindo/mCD8::GFP for cAMP measurements. For image analysis, regions of interest (ROIs) were selected within main cells or secondary cells, and fluorescence intensities were measured over successive time frames. For each ROI, the fluorescence intensity ratio at each time point (R) was normalized to the mean ratio during the baseline period (*R_0_*; −80 to −2 s), and the normalized ratio was expressed as relative fluorescence intensity (*R/R_0_*).

### *O*-propargyl puromycin (OPP) labeling

OPP labeling was performed with the Click-iT Plus OPP Alexa Fluor 488 Protein Synthesis Assay kit (Thermo Fisher Scientific C10456) according to the manufacturer’s guidelines with minor modifications. Male reproductive tracts were dissected and cultured in Schneider’s Drosophila medium. Dissected tissues were incubated for 6 hrs in Schneider’s Drosophila medium containing either serotonin, cycloheximide (CHX), or vehicle control. CHX (0.75 mM; Nacalai Tesque, 06741-91), a translational elongation inhibitor, was used as a negative control. Following the 6 hr incubation, tissues were transferred to fresh culture medium of identical composition, supplemented with 20 μM OPP, and incubated for an additional 30 min. Mock-treated samples incubated without OPP served as an additional negative control. Tissues were then rinsed once with 1× PBS and fixed in 4% formaldehyde for 20 min. Subsequent click-reaction and detection steps were performed according to the manufacturer’s protocol.

### Statistical analysis

All experiments were performed independently at least twice. Experiments were not randomized, and investigators were not blinded. All statistical analyses were performed using the “R” software environment.

## Data and materials availability

RNA-seq raw data have been deposited in DNA Data Bank of Japan (BioProject accession number PRJDB40147; Run accession numbers DRR900219-DRR900230). All other data are included in the manuscript and Supplementary Datasets S1 and S2.

## Acknowledgments

We thank Toshiro Aigaki, Olga V. Alekseyenko, Young-Joon Kim, Toshihiro Kitamoto, Satoru Kobayashi, Shu Kondo, Shigeyuki Koshikawa, Robert Maeda, Kyohei Mikami, Masayuki Miura, Akinao Nose, Jae H. Park, Nobuaki Tanaka, Jan A. Veenstra, Mariana F. Wolfner, Daisuke Yamamoto, the Kyoto Stock Center, the Kyorin Stock Center, the Bloomington Drosophila Stock Center, the Vienna Drosophila Resource Center, and the Developmental Studies Hybridoma Bank for providing us with fly stocks and other reagents. We are also grateful to Takehiro Kato, Toshiro Aigaki, Anne C. von Phillipsborn, Seong-Jin Kim, Young-Joon Kim, and Mariana F. Wolfner for helpful discussions, and to Eric Kubli for encouraging us to pursue this project. R.H. and To.A. were recipients of young scientist fellowships from the Japan Society for the Promotion of Science.

## Funding

Japan Society for the Promotion of Science grant 21J20279 (R.H.)

Japan Society for the Promotion of Science grant 15J00652 (To.A.)

Japan Society for the Promotion of Science grant 24K21960 (R.N.)

Program of the Joint Usage/Research Center for Developmental Medicine, IMEG, Kumamoto University grant K23-06 (R.N.)

Program of the Joint Usage/Research Center for Developmental Medicine, IMEG, Kumamoto University grant K24-15 (R.N.)

Ministry of Education, Culture, Sports, Science, and Technology, Promotion of Development of a Joint Usage/Research System Project: Coalition of Universities for Research Excellence Program (CURE) grant JPMXP1323015486 (A.N.)

Ministry of Education, Culture, Sports, Science, and Technology-Supported Program for the Inter-University Research Network for High-Depth Omics, IMEG, Kumamoto University (A.N.)

Takeda Science Foundation grant (Y.S.-N.)

Mitsubishi Foundation Natural Science Research grant (R.N.)

## Author contributions

Conceptualization: R.H., R.N.

Formal analysis: R.H., Y.K.

Funding acquisition: R.H., To.A., Y.S.-N., A.N., R.N.

Investigation: R.H., Y.K., H.-K.L., To.A., T.T., Ta.A.

Methodology: R.H., Y.K., R.T., Ts.A., A.K., A.N., Ta.A., R.N.

Project administration: R.H., R.N.

Resources: R.H., R.T., A.K., Ts.A., Y.S.-N., A.N., Ta.A., R.N.

Supervision: R.N.

Validation: R.H., Y.K.

Visualization: R.H., Y.K., Ta.A., R.N.

Writing—original draft: R.H., Y.K., Y.S.-N., Ta.A., R.N.

Writing—review & editing: R.H., R.T., H.-K.L., To.A., T.T., Ts.A., A.K., A.N., R.N.

