## Supplementary Figures S1-S15, and Table S1-S6 for "Sex peptide-SPR-serotonin signaling regulates mating-induced Sex peptide replenishment in *Drosophila*, suggesting an ancestral role in males"

### Supplementary Materials

#### Supplementary Figures

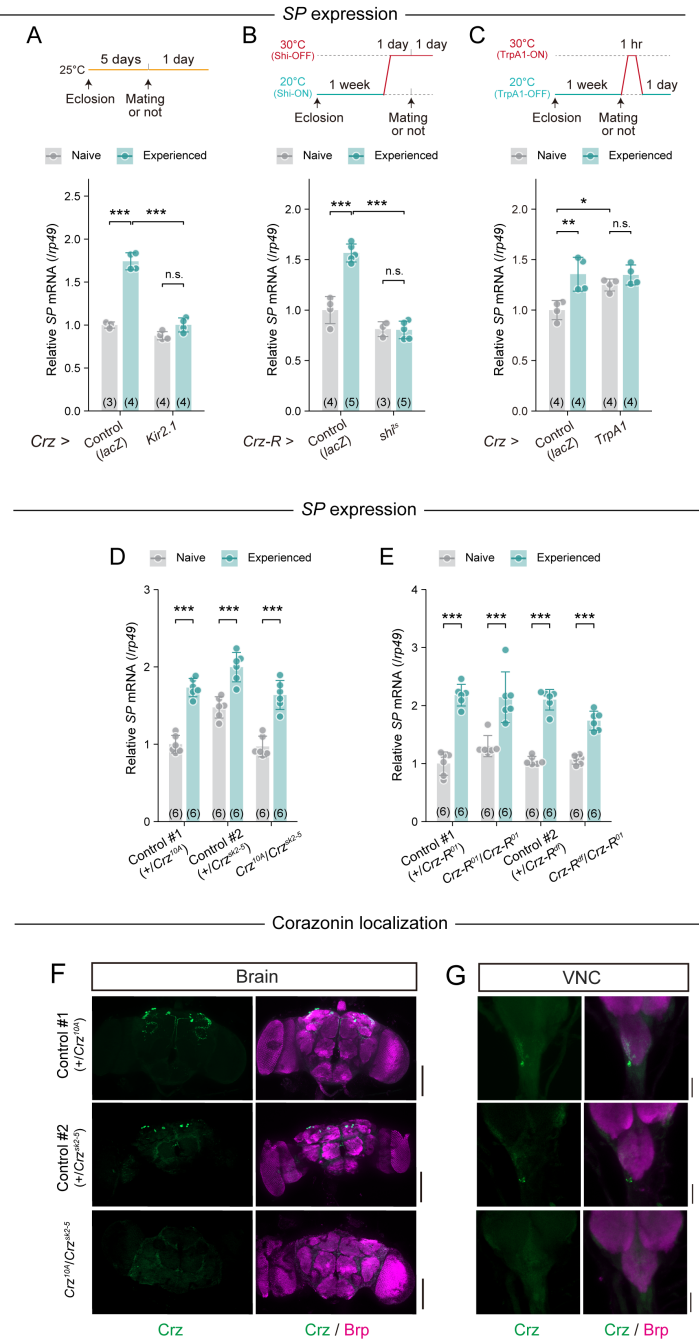

**Figure S1. Crz–Crz-R neurons are essential for mating-induced *SP* mRNA elevation but Crz–Crz-R signaling is dispensable.**

(A–E) *SP* mRNA levels before and after mating in male reproductive tracts from each indicated genotype. *SP* mRNA levels were measured by RT-qPCR. *SP* mRNA levels were calculated using *rp49* as the reference gene and subsequently normalized to the mean level in naive control males [*Crz* > *lacZ* in (A), *Crz-R* > *lacZ* in (B), *Crz* > *lacZ* in (C), +/*Crz<sup>10A</sup>* in (D), and +/*Crz-R<sup>01</sup>*

in (E)]. (A) *SP* mRNA levels in males with *Crz* neurons subjected to Kir2.1-mediated suppression (*Crz* > *Kir2.1*) and *lacZ* overexpression (*Crz* > *lacZ*) as a control. (B) *SP* mRNA levels in males subjected to *Crz*-R<sup>+</sup> neuron inhibition via *shi<sup>ts</sup>* (*Crz*-R > *shi<sup>ts</sup>*). These males were exposed to the restrictive temperature (30 °C) from one day before mating to dissection. *LacZ* overexpression (*Crz*-R > *lacZ*) was used as a control. (C) *SP* mRNA levels following activation of *Crz* neurons expressing *TrpA1* (*Crz* > *TrpA1*) at the restrictive temperature (30 °C) for 1 hr, one day before dissection. *lacZ* overexpression (*Crz* > *lacZ*) was used as a control. (D) *SP* mRNA levels in the *Crz* mutant (*Crz<sup>10A</sup>/Crz<sup>sk2-5</sup>*) and its controls (+/*Crz<sup>10A</sup>*, +/*Crz<sup>sk2-5</sup>*). (E) *SP* mRNA levels in *Crz*-R mutants (*Crz*-R<sup>01</sup>/*Crz*-R<sup>01</sup>, *Crz*-R<sup>01</sup>/*Crz*-R<sup>df</sup>) and their controls (+/*Crz*-R<sup>01</sup>, +/*Crz*-R<sup>df</sup>).

All quantitative data are presented as means ± SD. The number of samples (n) is indicated in each graph. Statistical analysis: Tukey-Kramer's HSD test for (A–E). \**P* < 0.05, \*\**P* < 0.01, \*\*\**P* < 0.001; n.s., non-significant (*P* > 0.05).

(F–G) Representative images of anti-corazonin (green) and anti-Brp (magenta) immunostaining of the (F) brain and (G) VNC of a *Crz* mutant (*Crz<sup>10A</sup>/Crz<sup>sk2-5</sup>*) and its controls (+/*Crz<sup>10A</sup>*, +/*Crz<sup>sk2-5</sup>*). Scale bars, (F) 100 μm, (G) 30 μm.

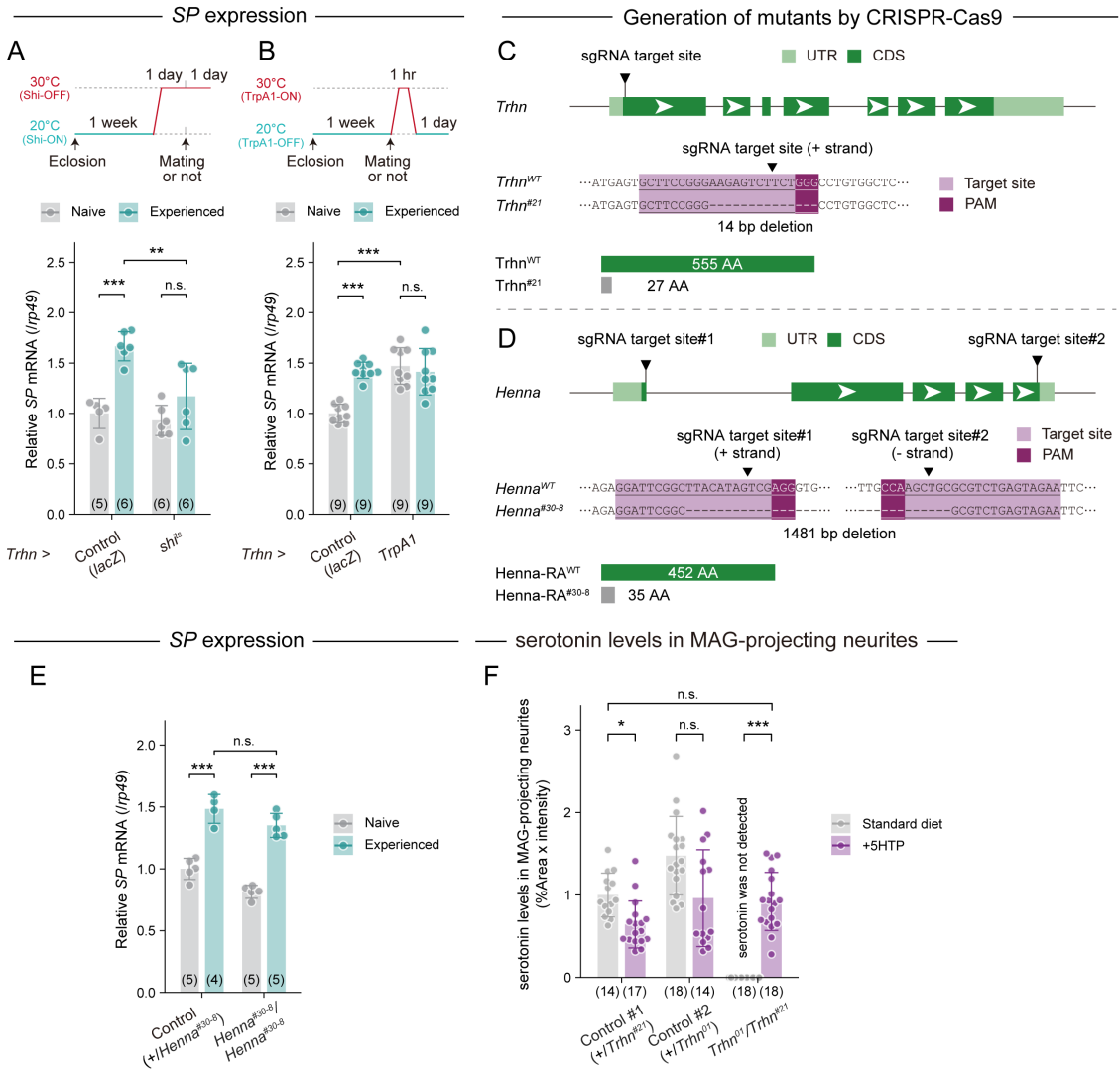

**Figure S2. Serotonergic neuron manipulation affects *SP* mRNA levels.**

(A, B, and E) *SP* mRNA levels before and after mating in male reproductive tracts of the indicated genotypes were measured via RT-qPCR. *SP* mRNA levels were calculated using *rp49* as the reference gene and subsequently normalized to the mean level in naive control males [*Trhn* > *lacZ* in (A), *Trhn* > *lacZ* in (B), and +/*Henna*<sup>#30-8</sup> in (E)]. (A) *SP* mRNA levels following serotonergic neuron inhibition via expression of *shi<sup>ts</sup>* (*Trhn* > *shi<sup>ts</sup>*) and exposure to the restrictive temperature (30 °C) from one day before mating to dissection. *LacZ* overexpression (*Trhn* > *lacZ*) was used as a control. (B) *SP* mRNA levels following serotonergic neuron activation via expression of *TrpA1* (*Trhn* > *TrpA1*) and exposure to the activating temperature (30 °C) for 1 hr, one day before dissection. *LacZ* overexpression (*Trhn* > *lacZ*) was used as a control. (C–D) Schematic indicating the sgRNA target sites (cleavage sites) in the (C) *Trhn* and (D) *Henna* genomic loci. CDS regions are dark green, while UTRs are light green. The target locus for each Cas9-induced mutant was PCR-amplified and sequenced. The WT sequence is shown at the top of each set of sequences as a reference. The Cas9-sgRNA target sequence is underlined, and the PAM indicated. Deleted nucleotides are indicated by dashes. Indel size is indicated next to each sequence. Each deletion causes a premature stop codon, resulting in truncated proteins (with some C-terminal amino acid extensions caused by frameshift) shown at the bottom of the panel. (E) *SP* mRNA levels in a *Henna* null mutant (*henna*<sup>#30-8</sup>/*henna*<sup>#30-8</sup>) and its control (+/*henna*<sup>#30-8</sup>). (F) Quantification of serotonin levels in the MAGs of *Trhn*-null mutant males (*Trhn*<sup>01</sup>/*Trhn*<sup>#21</sup>) and their corresponding controls (+/*Trhn*<sup>#21</sup>, +/*Trhn*<sup>01</sup>). Serotonin levels were measured using fluorescence intensity of anti-serotonin immunostaining in MAGs from males fed a standard diet with or without 5-hydroxytryptophan (5-HTP). Signal intensities were normalized to the mean signal intensity of control males with standard diet (+/*Trhn*<sup>#21</sup>). All quantitative data are presented as means ± SD. The number of samples (n) is indicated in each graph. Statistical analysis: Tukey-Kramer's HSD test for (A, B and E) and Steel-Dwass's multiple comparison test for (F). \*$P < 0.05$, \*\*$P < 0.01$, \*\*\*$P < 0.001$; n.s., non-significant ($P > 0.05$).

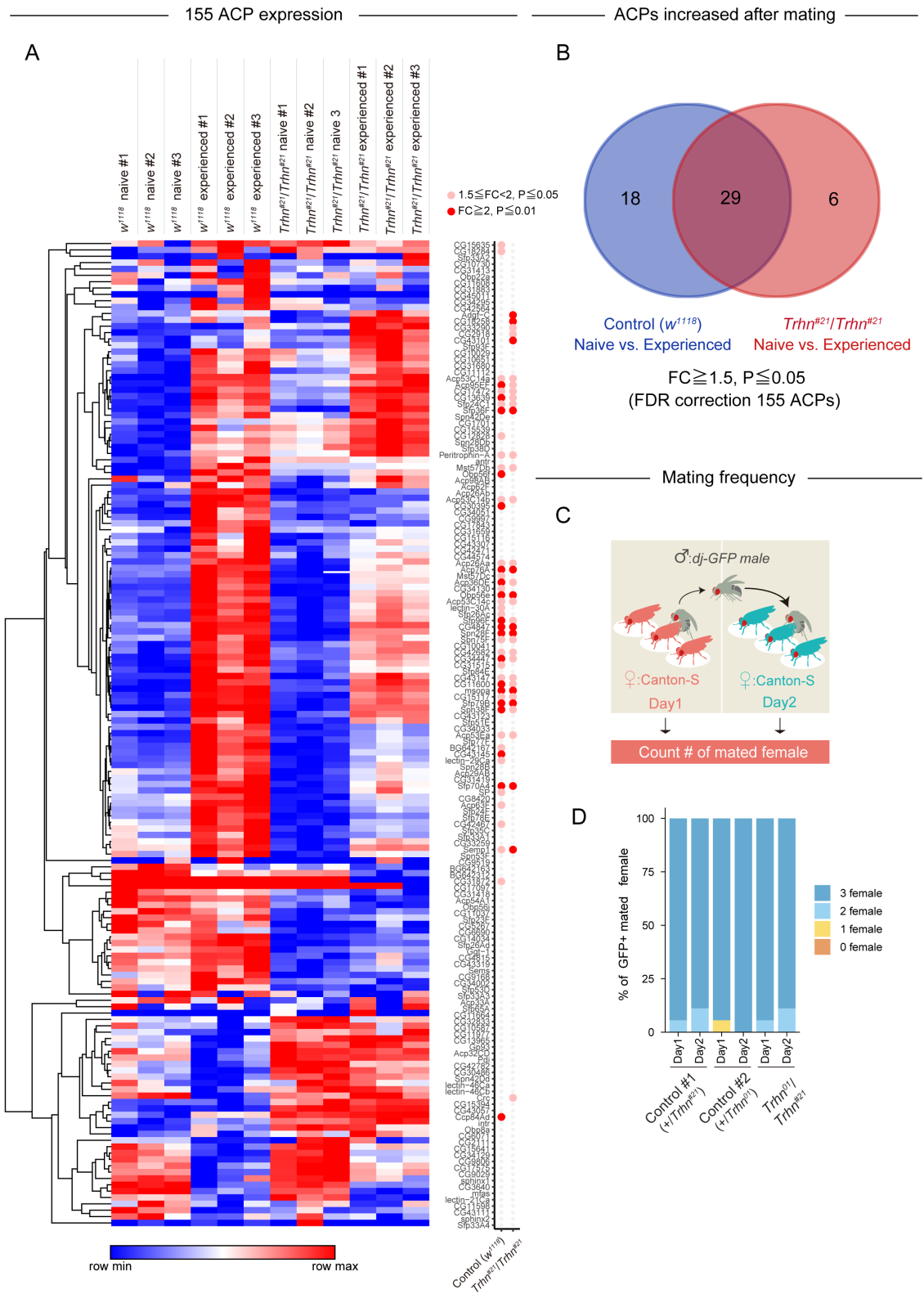

**Figure S3. *Trhn* is required for the post-mating increase in the expression of many accessory gland proteins (ACPs) beyond *SP*.**

(A) A heatmap showing RNA-seq-based expression profiles of 155 ACP genes in MAGs from $w^{1118}$ and *Trhn*-null mutant males ($Trhn^{#21}/Trhn^{#21}$) before and after mating. Colors in the heatmap represent $\log_2$-transformed normalized expression values. Dots displayed alongside the heatmap indicate the statistical significance and magnitude of post-mating upregulation, color-coded according to the following criteria: $FDR \leq 0.01$ and fold change (FC) $\geq 2$; $FDR \leq 0.05$ and $2 > FC \geq 1.5$ (B) Venn diagram showing the overlap of genes in $w^{1118}$ and *Trhn*-null mutant males upregulated after mating. Upregulated genes were defined as those with $FDR \leq 0.05$ and $FC \geq 1.5$. (C) Schematic showing the protocol for examining mating frequency using the same mating protocol as described in the legend for Fig. 2H. (D) Mating frequencies for each group with the labels indicating the presence of green fluorescent sperm in 0, 1, 2, or 3 females. The number of samples (n) is indicated above the graph.

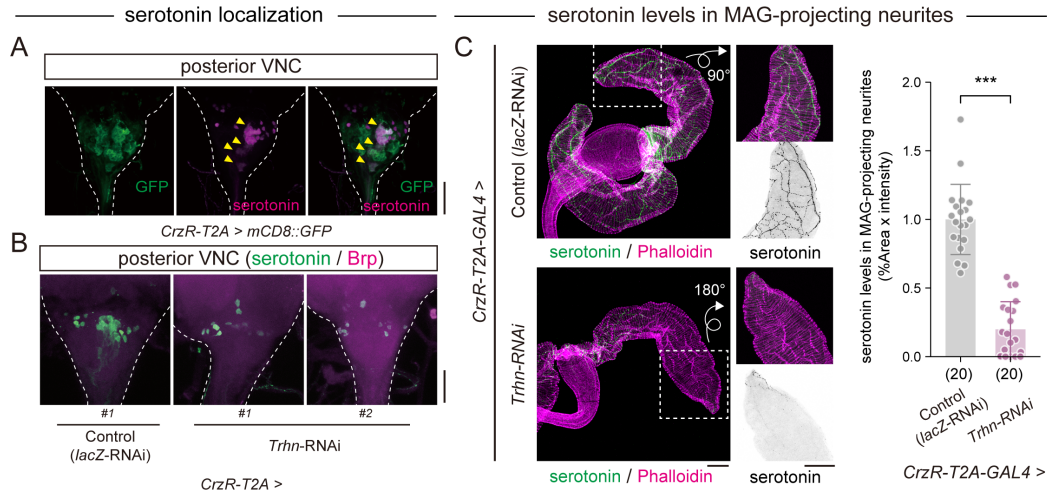

**Figure S4. *CrzR-GAL4* effectively labels serotonergic neurons projecting to the MAGs.**

(A) *CrzR-T2A-GAL4*-driven mCD8::GFP (green) and anti-serotonin (magenta) immunohistochemistry in the posterior VNC. Scale bar, 30 μm. (B) Representative images of anti-serotonin (green) and anti-Brp (pan-neuronal marker; magenta) immunohistochemistry in the posterior VNC from *Trhn* knockdown males. Scale bar, 30 μm. (C) Left panel: Representative images of male reproductive tracts from naive *Trhn* knockdown males (*CrzR > Trhn-RNAi*) and their controls (*CrzR > lacZ-RNAi*) stained with an anti-serotonin antibody (green) and phalloidin (magenta). Scale bar, 100 μm. Right panel: Serotonin levels in the MAG of *Trhn* knockdown males (*CrzR > Trhn-RNAi*) and their controls. Serotonin levels were normalized to the mean values of control males. All quantitative data are presented as means ± SD. The number of samples (n) is indicated in each graph. Statistical analysis: Wilcoxon rank sum test for (C). \*\*\*$P < 0.001$; n.s., non-significant ($P > 0.05$).

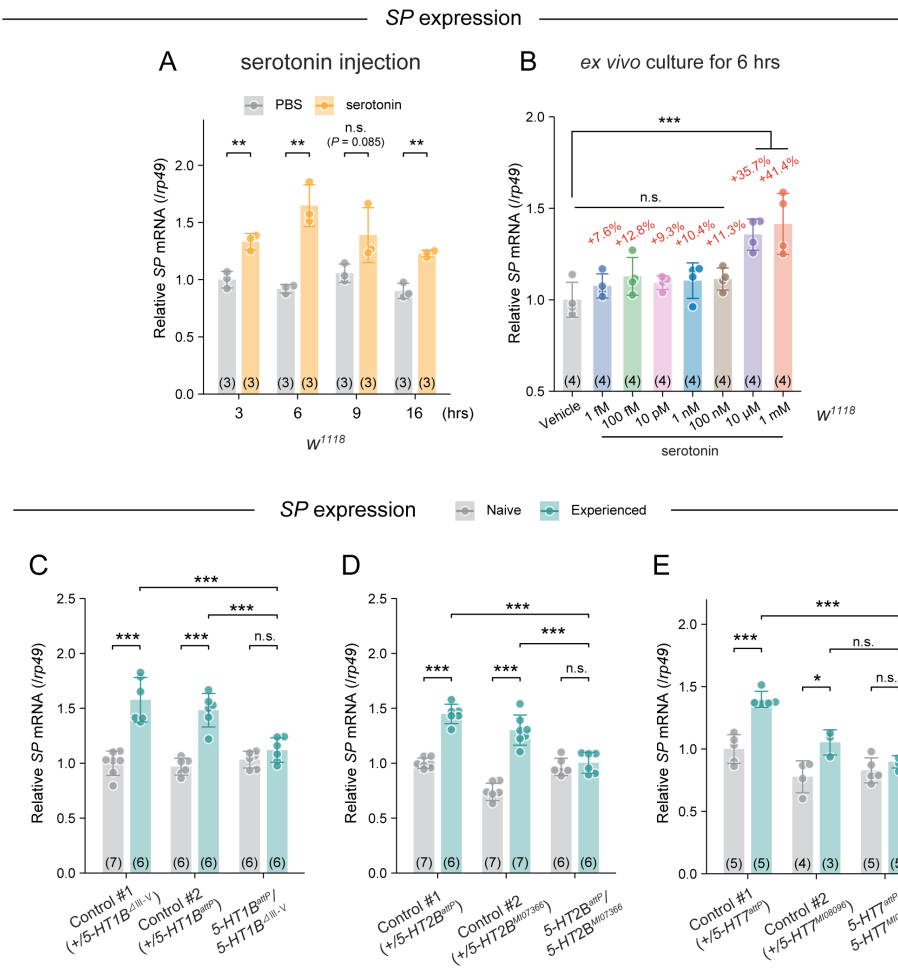

**Figure S5. Involvement of serotonin and its receptors in *SP* mRNA expression in the MAG.**

(A–E) *SP* mRNA levels in male reproductive tracts of the indicated genotypes were measured via RT-qPCR. *SP* mRNA levels were calculated using *rp49* as the reference gene and subsequently normalized to the mean level in naive control males [PBS at 3 h in (A), vehicle in (B), +/5-*HT1B*<sup>III-V</sup> in (C), +/5-*HT2B*<sup>attP</sup> in (D), and +/5-*HT7*<sup>attP</sup> in (E)]. (A) *SP* mRNA levels in naive *w<sup>1118</sup>* males 3, 6, 9, and 16 hrs after injection of PBS (control) or 1 mM serotonin. (B) *SP* mRNA levels in male reproductive tracts of naive males cultured *ex vivo* for 6 hrs with 1 fM – 1 mM serotonin. (C–E) The 5-HT1B, 5-HT2B, and 5-HT7 serotonin reception promote *SP* mRNA upregulation in MAGs. *SP* mRNA levels before and after mating in the male reproductive tracts of the indicated genotypes. *SP* mRNA levels were measured via RT-qPCR and normalized to *rp49*. Values are presented as means ± SD. (C) *SP* mRNA levels in 5-*HT1B* mutant (5-*HT1B*<sup>III-V</sup>/5-*HT1B*<sup>attP</sup>) and control animals (#1: +/5-*HT1B*<sup>III-V</sup>, #2: +/5-*HT1B*<sup>attP</sup>). (D) *SP* mRNA levels in 5-*HT2B* mutant (5-*HT2B*<sup>M107366</sup>/5-*HT2B*<sup>attP</sup>) and control animals (#1: +/5-*HT2B*<sup>attP</sup>, #2: +/5-*HT2B*<sup>M107366</sup>). (E) *SP* mRNA levels in 5-*HT7* mutant (5-*HT7*<sup>attP</sup>/5-*HT7*<sup>M108096</sup>) and control animals (#1: +/5-*HT7*<sup>M108096</sup>, #2: +/5-*HT7*<sup>attP</sup>). All quantitative data are presented as means ± SD. The number of samples (n) is indicated in each graph. Statistical analysis: unpaired two-tailed Student's *t*-test for (A); Dunnett's test for (B); Tukey-Kramer's HSD test for (C–E).
\*$P < 0.05$, \*\*$P < 0.01$, \*\*\*$P < 0.001$; n.s., non-significant ($P > 0.05$).

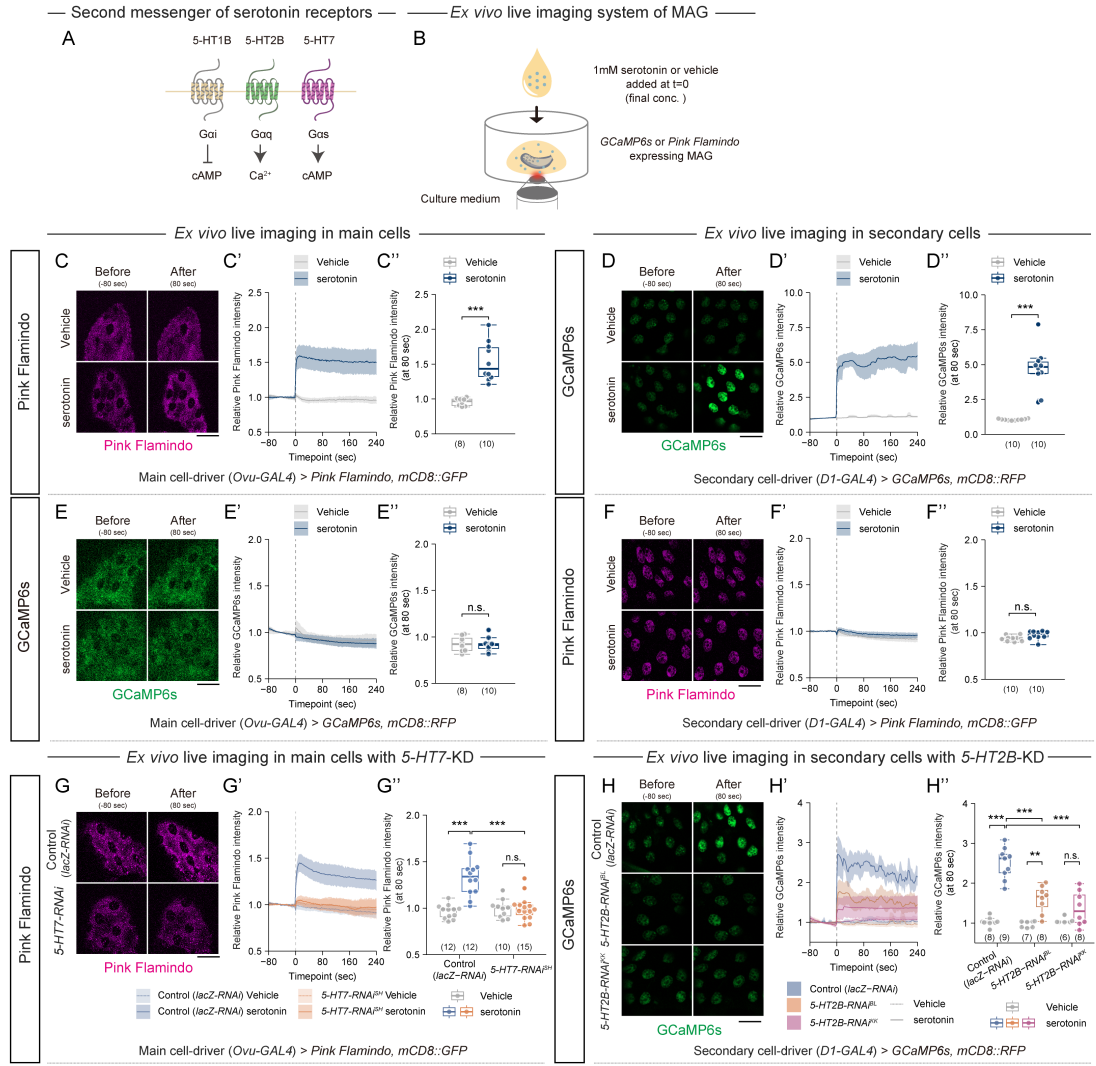

**Figure S6. Serotonin increases intracellular Ca<sup>2+</sup> and cAMP levels in the MAG via serotonin receptors.**

(A) Schematic showing the second messengers downstream of each serotonin receptor. (B) Schematic for the live imaging of MAGs cultured *ex vivo* with serotonin. (C–H) Pink Flamindo and GCaMP6s fluorescence reporters were expressed in main and secondary cells to permit live imaging of second messenger activity. (C–F) Control (wild-type) genetic background. (G) 5-HT7 knockdown in main cells. (H) 5-HT2B knockdown in secondary cells. Left panel: Representative images of Pink Flamindo (magenta) or GCaMP6s (green) in main cells or secondary cells before (-80 sec) and after (80 sec) serotonin administration. Scale bars, 50 μm. Middle panel ('): Temporal changes in fluorescence intensity from reporters in main cells or secondary cells. Medium, with or without the addition of serotonin, was applied at t=0. Pink Flamindo fluorescence was normalized to mCD8::GFP, and GCaMP6s fluorescence was normalized to mCD8::RFP. Values are shown as means with 95% confidence intervals. Right panel (''): Relative intensity at t = 80 was calculated by normalized intensity from t = -80 to -2 (set to 1.0). Box-and-whisker plots showing the normalized Pink Flamindo or GCaMP6s signals in main cells or secondary cells in response to serotonin. The number of samples (n) is indicated in each graph. All scale bars, 50 μm. Statistical analysis: unpaired two-tailed Student's *t*-test for

(C'', D'', E'', and F'') and Tukey-Kramer's HSD test for (G'' and H''). \*$P < 0.05$, \*\*$P < 0.01$, \*\*\*$P < 0.001$; n.s., non-significant ($P > 0.05$).

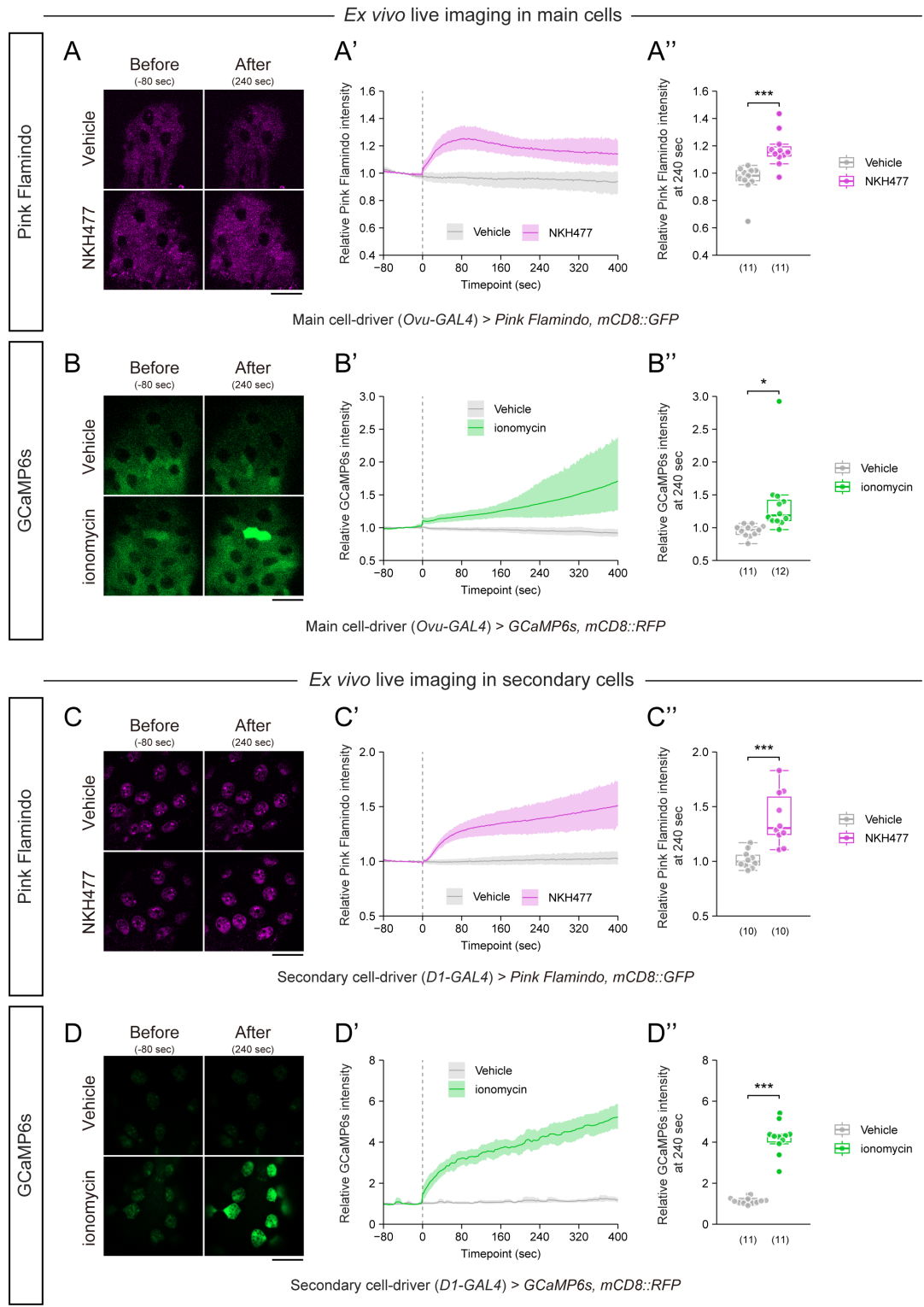

**Figure S7. Ionomycin and NKH477 increase intracellular $\text{Ca}^{2+}$ and cAMP levels in the MAG, respectively.**

(A–D) PinkFlamindo and GCaMP6s were expressed in main cells and secondary cells to permit live imaging of second messengers. Left panels: Representative images of Pink Flamindo (magenta) or GCaMP6s (green) in main cells and secondary cells before (-80 sec) and after (240 sec) addition of NKH477 or ionomycin. Scale bars, 50 μm. Middle panels (''); Temporal changes in reporter fluorescence in main cells or secondary cells with or without NKH477 or ionomycin. Pink Flamindo fluorescence was normalized to mCD8::GFP, and GCaMP6s fluorescence was normalized to mCD8::RFP. Medium, with or without NKH477 or ionomycin, was applied at $t = 0$. Values are shown as means with 95% confidence intervals. Right panels (''); Relative intensity at $t = 240$ was calculated by normalized intensity from $t = -80$ to $-2$ (set to 1.0). Box-and-whisker plots showing normalized Pink Flamindo or GCaMP6s fluorescence in main cells or secondary cells after addition of NKH477 or ionomycin. (A) *Ex vivo* live imaging of Pink Flamindo in main cells with or without NKH477. (B) *Ex vivo* live imaging of GCaMP6s in main cells with or without ionomycin. (C) *Ex vivo* live imaging of Pink Flamindo in secondary cells with or without NKH477. (D) *Ex vivo* live imaging of GCaMP6s in secondary cells with or without ionomycin. The number of samples (n) is indicated in each graph. All scale bars: 50 μm. Statistical analysis: unpaired two-tailed Student's *t*-test. \*$P < 0.05$, \*\*$P < 0.01$, \*\*\*$P < 0.001$; n.s., non-significant ($P > 0.05$).

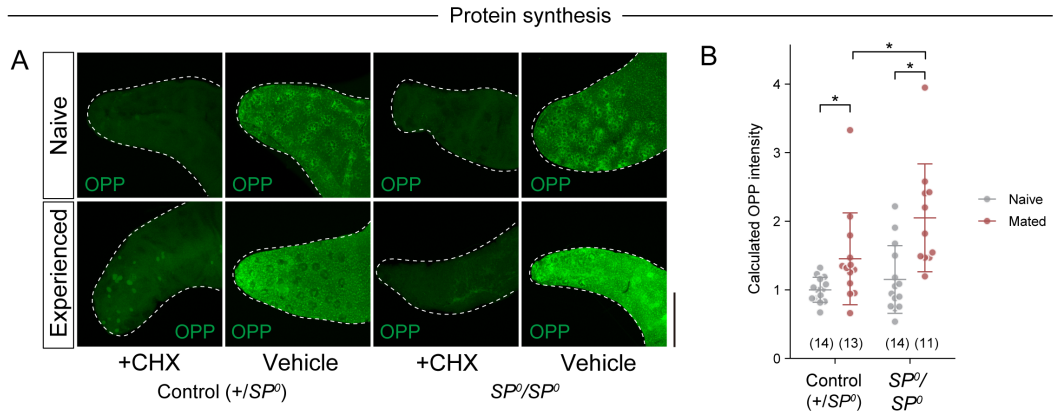

**Figure S8. OPP staining in MAGs of naive and experienced control and $SP^0$ mutant males ($SP^0/SP^0$).**

(A–B) Dissected male reproductive tracts were cultured with OPP to visualize *de novo* protein synthesis in the MAGs. (A) Representative images showing OPP fluorescence (green) in MAGs (outlined by dashed lines) before and after mating, with or without cycloheximide (CHX) treatment. Scale bar, 100 μm. (B) Quantification of OPP fluorescence intensity in MAGs before and after mating. Signal intensities were normalized to the mean signal intensity of naive control males (+/SP<sup>0</sup>). All quantitative data are presented as means ± SD. The number of samples (n) is indicated in each graph. Statistical analysis: Steel–Dwass multiple comparisons test. \*$P < 0.05$, ; n.s., non-significant ($P > 0.05$).

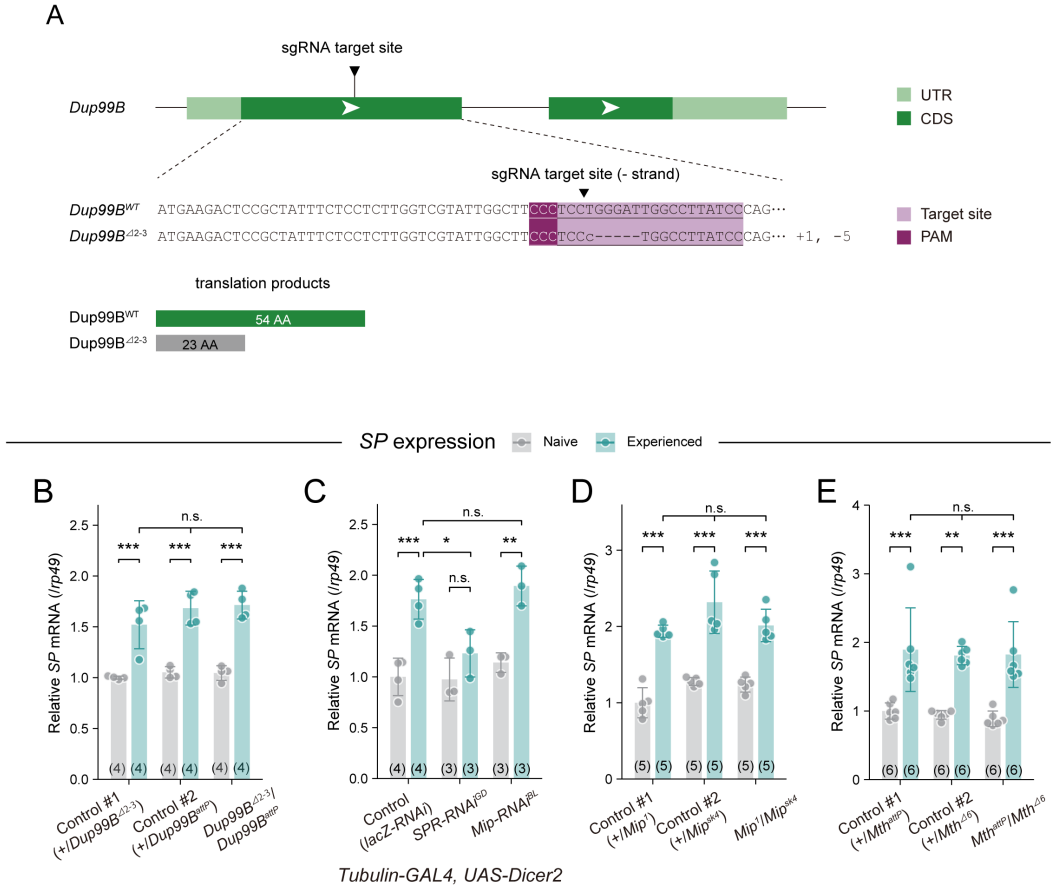

**Figure S9. *Dup99B*, *Mip*, and *Mth* are unnecessary for the post-mating increase in *SP* mRNA.**

(A) Schematic showing the sgRNA target sites (cleavage sites) in the *Dup99B* genomic locus. CDS regions are dark green, and introns are light green. The target locus in each Cas9-induced mutant was PCR-amplified and sequenced. The WT sequence is shown at the top of each set of sequences as a reference. The Cas9-sgRNA target sequence is underlined, and the PAM is indicated. Deleted nucleotides are shown as dashes. Indel size is indicated next to each sequence. Indel mutations result in premature stop codons. (B–E) *SP* mRNA levels before and after mating in the male reproductive tract of the indicated genotypes. *SP* mRNA levels were measured by RT-qPCR. *SP* mRNA levels were calculated using *rp49* as the reference gene and subsequently normalized to the mean level in naive control males [*+Dup99B*<sup>Δ2-3</sup> in (B), *Tubulin-GAL4* > *UAS-Dicer lacZ-RNAi* in (C), *+Mip*<sup>1</sup> in (D), and *+Mth*<sup>attP</sup> in (E)]. Values are presented as means ± SD. (B) *SP* mRNA levels in *Dup99B* mutant (*Dup99B*<sup>Δ2-3</sup>/*Dup99B*<sup>attP</sup>) and control animals (#1: *+Dup99B*<sup>Δ2-3</sup> #2: *+Dup99B*<sup>attP</sup>). (C) *SP* mRNA levels in males with systemic knockdown of *SPR* (*SPR-RNAi*), and *Mip* (*Mip-RNAi*) under the control of *Tubulin-GAL4*. *LacZ* knockdown (*lacZ-RNAi*) was used as a control. (D) *SP* mRNA levels in *Mip* mutant (*Mip*<sup>01</sup>/*Mip*<sup>sk4</sup>) and control animals (#1: *+Mip*<sup>01</sup> #2: *+Mip*<sup>sk4</sup>). (E) *SP* mRNA levels in *Mth* mutant (*Mth*<sup>attP</sup>/*Mth*<sup>Δ6</sup>) and control animals (#1: *+Mth*<sup>attP</sup> #2: *+Mth*<sup>Δ6</sup>). All quantitative data are presented as means ± SD. The number of samples (n) is indicated in each graph. Statistical

analysis: Tukey-Kramer's HSD test for (B–E). \*$P < 0.05$, \*\*$P < 0.01$, \*\*\*$P < 0.001$; n.s., non-significant ($P > 0.05$).

A

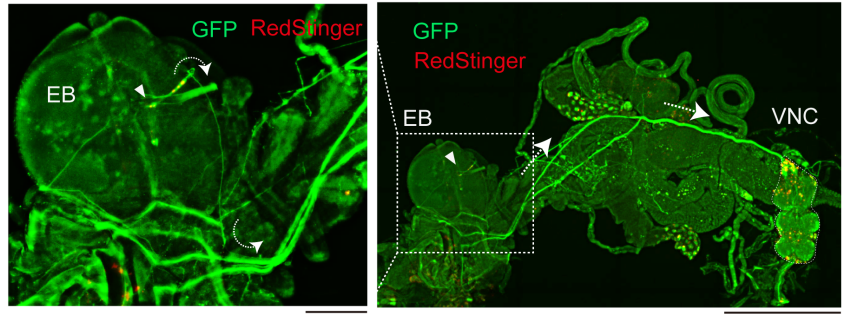

*R40A05-GAL4 > UAS-myr::GFP-T2A-RedStinger*

B

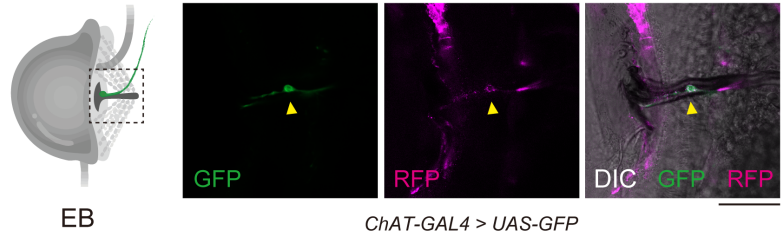

*ChAT-GAL4 > UAS-GFP*
*SPR-T2A-LexA > LexAop-CD4-td-Tomato*

C

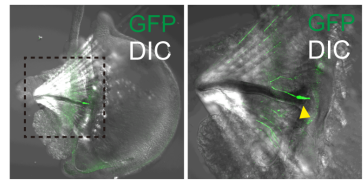

*SPR-T2A-GAL4;*
*UAS > stop > GFP; fru-Flp*

D

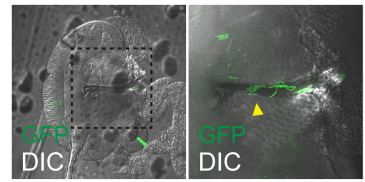

*SPR-T2A-GAL4;*
*UAS > stop > GFP; dsx-Flp*

**Figure S10. *SPR*<sup>+</sup> EB neurons share characteristics with female SPSNs.**

(A) Localization of a membrane-tethered GFP (green) and the nuclear marker RedStinger (red) under the control of the *R40A05*-GAL4 driver. The left panel shows the area around the EB. White arrowheads indicate cell bodies. The right panel shows a lower magnification view of a neuron connecting the EB and VNC. Scale bars, left: 100 μm, right: 500 μm. (B) Visualizing *ChAT-GAL4* driving GFP (green) and *SPR-T2A-LexA* driving mTd-Tomato (magenta) in the EB. The right image is an overlap of the GFP and mTd-Tomato channels with the differential interferogram. Yellow arrowheads indicate a *GFP*<sup>+</sup> *RFP*<sup>+</sup> cell body. Scale bar: 50 μm. (C–D) Representative images showing GFP reporter expression in (C) *fru*<sup>+</sup> *SPR*<sup>+</sup> neurons or (D) *dsx*<sup>+</sup> *SPR*<sup>+</sup> neurons from flies carrying *SPR-T2A-GAL4* with *fru-FLP* or *dsx-FLP*. GFP fluorescence has been superimposed on a differential interference contrast (DIC) image. The left panel shows the EB, while the right panel shows a higher-magnification view of the region in the left image.

enclosed by the dashed box. The yellow arrowhead indicates a GFP-labeled neuron. Scale bar: 50 $\mu\text{m}$.

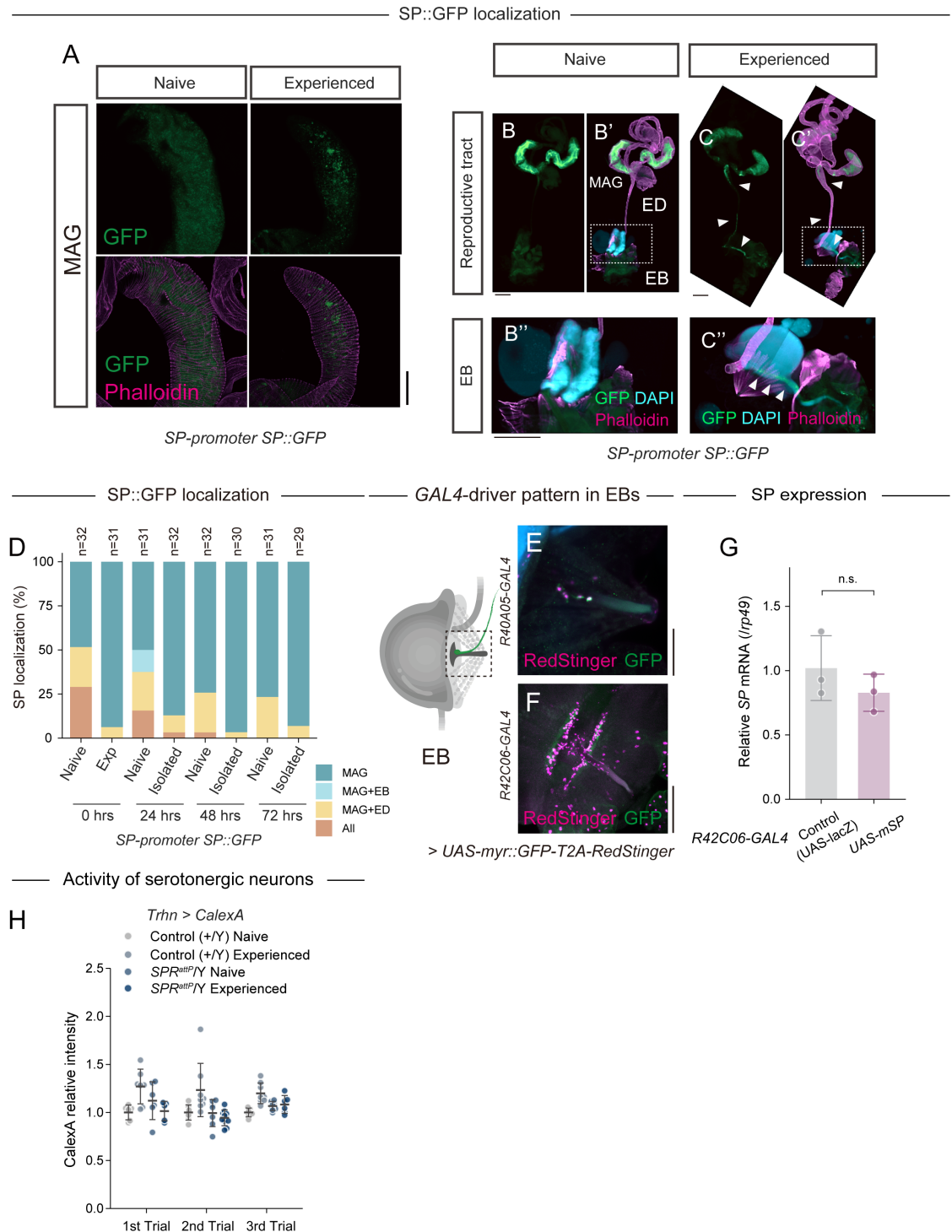

**Figure S11. SP localization and neuronal analyses related to post-mating SP replenishment.**

(A–C) Localization of SP::GFP in the male reproductive tract before and after mating. (A) Representative images of SP::GFP (green) and phalloidin (magenta) staining in the male reproductive tract before and after mating. Scale bar, 100 μm. (B and C) Representative images of SP::GFP (green), phalloidin (magenta), and DAPI (cyan) staining in the male reproductive tract before and after mating. Scale bar, 200 μm. White arrowheads indicate SP::GFP in the ED or EB. Scale bar, 200 μm. (D) Temporal changes in SP-GFP localization in the male reproductive tract of *SP-promoter-SP::GFP* (*SP::GFP*) lines after mating. SP::GFP was localized to the MAG only, the MAG and EB, the MAG and ED, or the MAG and EB and ED. (E–F) Representative images of myr::GFP (green) and RedStinger (magenta) in the EB under the control of the (E) *R40A05-GAL4* and (F) *R42C06-GAL4* drivers. Scale bar, 50 μm. Yellow arrowheads indicate cell bodies. (G) RT-qPCR analysis of *SP* mRNA levels in the male reproductive tracts of control (*R42C06-GAL4 > UAS-lacZ*) and membrane-tethered SP (*UAS-mSP: R42C06-GAL4 > UAS-mSP*) males. *SP* mRNA levels were calculated using *rp49* as the reference gene and subsequently normalized to the mean level in control males. (H) Original data showing CaLexA in serotonergic neurons (*Trhn > CaLexA*) located in the posterior VNC of *SPR* mutant males (*SPR<sup>attP</sup>/Y*) and their controls (+/Y) before and after mating. Signal intensities were normalized in each trial to the mean of the control naive males. Related to Figure 4, L–M. All quantitative data are presented as means ± SD. The number of samples (n) is indicated in each graph. Statistical analysis: unpaired two-tailed Student's *t*-test for (G). \**P* < 0.05, \*\**P* < 0.01, \*\*\**P* < 0.001; n.s., non-significant (*P* > 0.05).

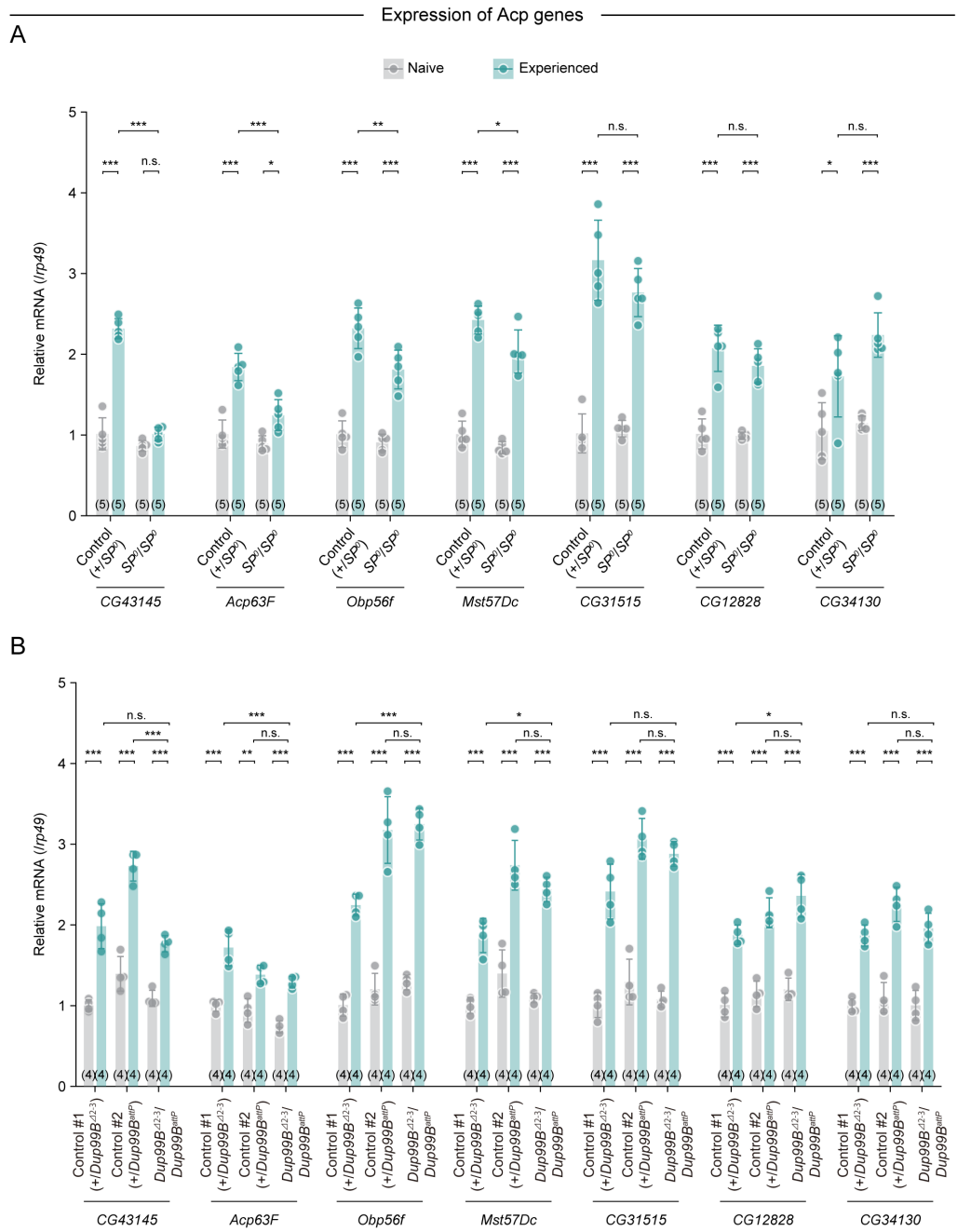

**Figure S12. *SP* is necessary for the post-mating increase in several Acp mRNAs.**

(A) RT-qPCR analysis of Acp mRNA levels in the male reproductive tracts of control (+/*SP*<sup>0</sup>) and *SP* knockout mutant (*SP*<sup>0</sup>/*SP*<sup>0</sup>) males. For each transcript, mRNA levels were calculated using *rp49* as the reference gene and subsequently normalized to the mean level in naive control males. (B) RT-qPCR analysis of Acp mRNA levels in the male reproductive tracts of control #1 (+/*Dup99B*<sup>Δ2-3</sup>), control #2 (+/*Dup99B*<sup>attP</sup>) and *Dup99B* mutant (*Dup99B*<sup>Δ2-3</sup>/*Dup99B*<sup>attP</sup>). For each transcript, mRNA levels were calculated using *rp49* as the reference gene and subsequently normalized to the mean level in naive control #1 males. All quantitative data are presented as means ± SD. The number of samples (n) is indicated in each graph. Statistical analysis: Tukey-Kramer's HSD test. \*$P < 0.05$, \*\*$P < 0.01$, \*\*\*$P < 0.001$; n.s., non-significant ($P > 0.05$).

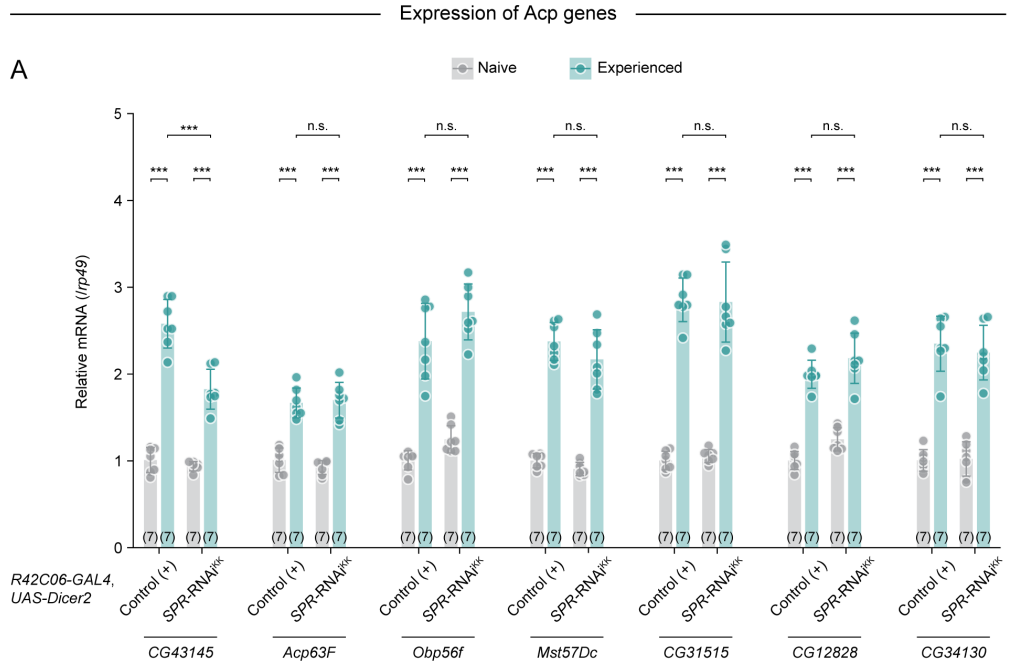

**Figure S13. *SPR* knockdown in *R42C06-GAL4*-positive neurons reduces post-mating *CG43145* mRNA levels.**

(A) RT-qPCR analysis of Acp mRNA levels in the male reproductive tracts of control (*R42C06-GAL4* > *UAS-Dicer2*) and *SPR-RNAi* (*R42C06-GAL4* > *UAS-Dicer2*, *SPR-RNAi*<sup>KK</sup>) males. For each transcript, mRNA levels were calculated using *rp49* as the reference gene and subsequently normalized to the mean level in naive control males. All quantitative data are presented as means ± SD. The number of samples (n) is indicated in each graph. Statistical analysis: Tukey-Kramer's HSD test. \*$P$ < 0.05, \*\*$P$ < 0.01, \*\*\*$P$ < 0.001; n.s., non-significant ($P$ > 0.05).

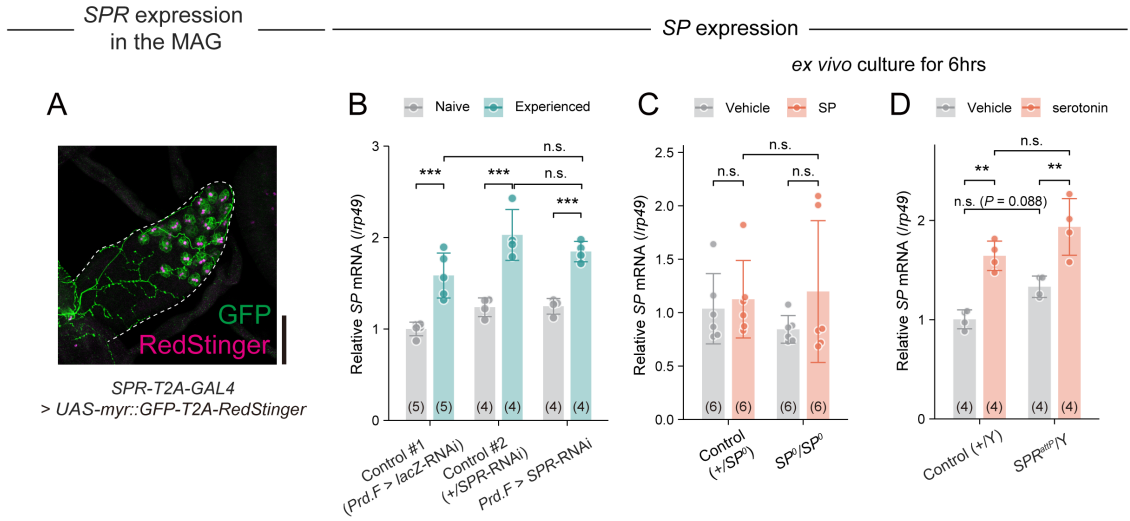

**Figure S14. SP seems not to act on the MAG directly for post-mating SP replenishment.** (A) Representative image of *SPR-T2A-GAL4* driving myr::GFP (green) and RedStinger (magenta) in the MAG of wild-type males. Scale bar, 50 μm. (B) *SP* mRNA levels before and after mating in the male reproductive tracts of males with *Prd.F-GAL4* driving *SPR* RNAi (*Prd.F > SPR-RNAi*) and their controls (*Prd.F > lacZ-RNAi*, *SPR-RNAi/+*). *SP* mRNA levels were determined by RT-qPCR. *SP* mRNA levels were calculated using *rp49* as the reference gene and subsequently normalized to the mean level in control males (*Prd.F > lacZ-RNAi*). (C) *SP* mRNA levels in *ex vivo*-cultured male reproductive tracts of naive control (+/*SP<sup>0</sup>*) and *SP<sup>0</sup>* males (*SP<sup>0</sup>/SP<sup>0</sup>*) with or without 1 μM SP. Male reproductive tracts were treated with SP for 6 hrs. Samples were measured by RT-qPCR. *SP* mRNA levels were calculated using *rp49* as the reference gene and subsequently normalized to the mean level in control males. (D) *SP* mRNA levels in the male reproductive tracts of naive *SPR* mutant (*SPR<sup>attP</sup>/Y*) males and their controls (+/*Y*) after 6 hrs of *ex vivo* culture with or without 1 mM serotonin. Samples were measured via RT-qPCR. *SP* mRNA levels were calculated using *rp49* as the reference gene and subsequently normalized to the mean level in control males. All quantitative data are presented as means ± SD. The number of samples (n) is indicated in each graph. Statistical analysis: Tukey-Kramer's HSD test for (B, C and D). \*$P < 0.05$, \*\*$P < 0.01$, \*\*\*$P < 0.001$; n.s., non-significant ($P > 0.05$).

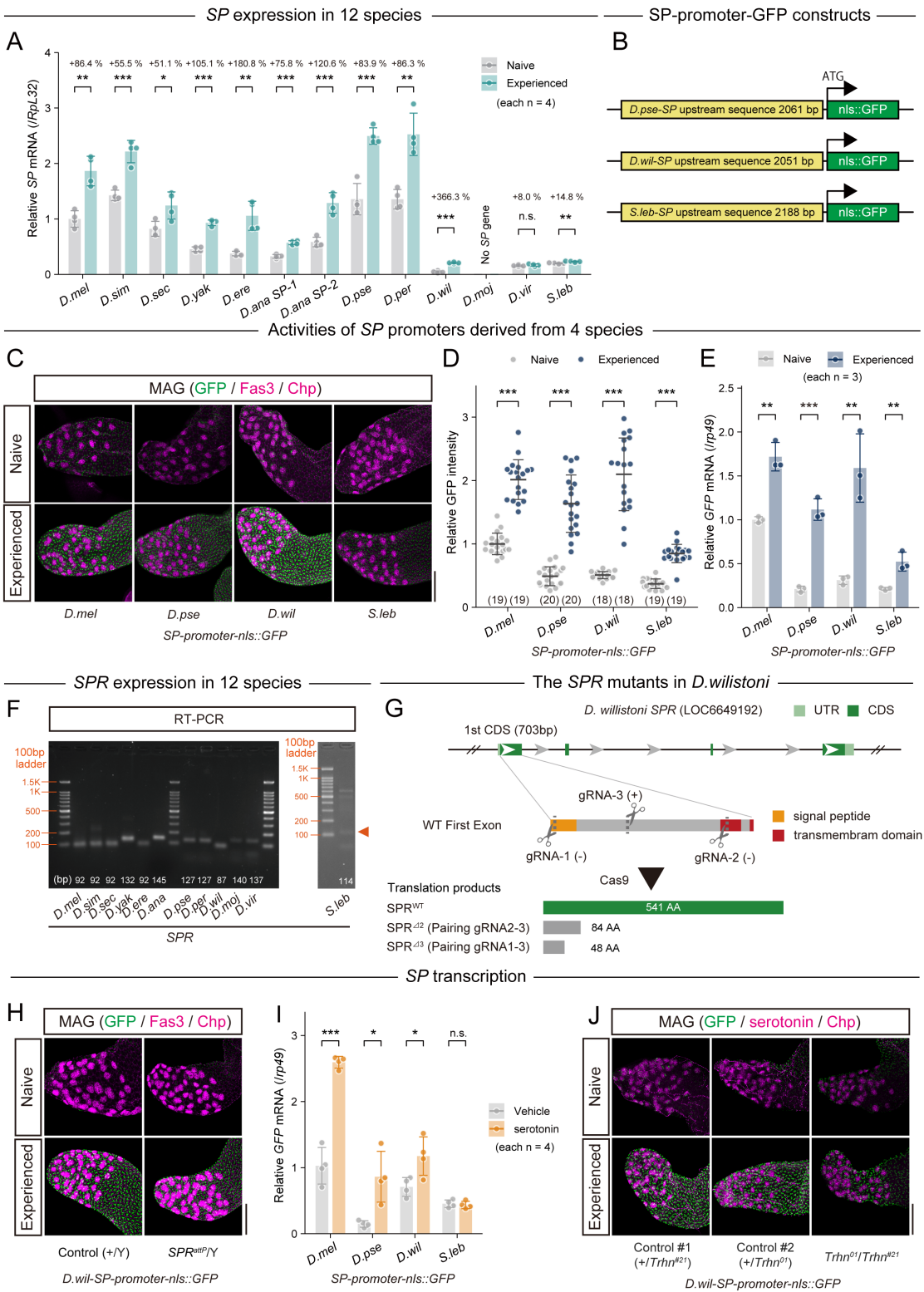

**Figure S15. Comparative analyses of *SP* expression, *SP*-promoter activity, and *SPR* function in drosophilid males.**

Abbreviations: *D. melanogaster*, *D. mel*; *D. simulans*, *D. sim*; *D. sechellia*, *D. sec*; *D. yakuba*, *D. yak*; *D. erecta*, *D. ere*; *D. ananassae*, *D. ana*; *D. pseudoobscura*, *D. pse*; *D. persimilis*, *D. per*; *D. willistoni*, *D. wil*; *D. mojavensis*, *D. moj*; *D. virilis*, *D. vir*; and *S. lebanonensis*, *S. leb*.

(A) Relative *SP* mRNA levels before and after mating in the male reproductive tracts of 12 species, excluding only *D. moj*. Values are shown relative to the levels in naive *D. mel*. Samples were normalized to *rp49* (for *D. melanogaster*) and *RpL32* (for other *Drosophila* species). *SP* mRNA from each species and *RpL32* mRNA levels were quantified with standard curves made using a plasmid containing fragments of *SP* and *RpL32*. (B) Sequences upstream of the *SP* genes in *D. pse*, *D. wil*, and *S. leb* were used as *SP*-promoters. Genomic sequences upstream of the *SP* CDS were amplified and cloned to make the *SP*-promoter constructs. (C–E) *SP*-promoter activity before and after mating in *D. melanogaster* males carrying *GFP* reporter transgenes driven by *SP* promoters from *D. melanogaster*, *D. pseudoobscura*, *D. willistoni*, or *S. lebanonensis*. (C) Representative images of MAGs immunostained with anti-GFP (green), anti-Fas3 (magenta), and anti-Chp (magenta) antibodies. Scale bar, 100 μm. Promoter activity was assessed by quantifying GFP fluorescence intensity in immunostained MAGs (D) and *GFP* mRNA levels by RT-qPCR using *rp49* as the reference gene (E). Values in (D) and (E) were normalized to the corresponding mean values in naive males carrying the *D. melanogaster SP*-promoter reporter. (F) *SPR* gene expression in the male reproductive tract of 12 species as measured by RT-PCR. The numbers at the bottom of the image indicate amplicon length. (G) Schematic showing the sgRNA target sites (cleavage sites) in the *D.wil-SPR* genomic locus (LOC6649192). Three gRNAs targeting the first exon were designed and used to generate two deletion mutants lacking either part of the signal peptide or the transmembrane domain. The target locus in each Cas9-induced mutant was PCR-amplified and sequenced. The WT sequence is shown at the top of each set of sequences as a reference. Indel mutations caused a premature stop codon, resulting in truncated proteins (with some C-terminal amino acid extension caused by frameshift). (H) *D. wil-SP*-promoter activity before and after mating in the *SPR* mutant (*SPR<sup>attP</sup>/Y*) and its control (+/Y). Representative images of anti-GFP (green), anti-Fas3 (main cells; magenta), and anti-Chp (secondary cells; magenta) signal in the MAG. Scale bar, 100 μm. (I) *D.mel*-, *D.pse*-, *D.wil*-, and *S.leb-SP*-promoter activity in male reproductive tracts after 6 hrs of ex vivo culture with or without 1 mM serotonin. Promoter activity was determined by quantifying *GFP* mRNA levels via RT-qPCR. The values were calculated using *rp49* as the reference gene and subsequently normalized to the mean level in *D.mel*. (J) *D. wil-SP*-promoter activity before and after mating in the *Trhn* mutant (*Trhn<sup>01</sup>/Trhn<sup>#21</sup>*) and its controls (+/*Trhn<sup>01</sup>*, +/*Trhn<sup>#21</sup>*). Left panels: Representative images of anti-GFP (green), anti-Chp (magenta), and anti-serotonin (magenta) signals in the MAG. Scale bar, 100 μm. For C, H, and J, note that the most prominent magenta signals in the apical MAG correspond to secondary cells, which are anti-Chp-positive. All quantitative data are presented as means ± SD. The number of samples (n) is indicated in each graph. Statistical analysis: unpaired two-tailed Student's *t*-test for (A, E, and I) and Wilcoxon rank sum test for (D). \**P* < 0.05, \*\**P* < 0.01, \*\*\**P* < 0.001; n.s., non-significant (*P* > 0.05).

#### Supplementary Tables

**Table S1.** List of fly strains used in this study

*Drosophila melanogaster* and *Drosophila willistoni* strains

| Genotype | Source | Identifier | Stock# |
| --- | --- | --- | --- |
| Canton-S |  |  |  |
| *w<sup>1118</sup>* |  |  |  |
| *Crz-GAL4* | Bloomington Drosophila Stock Center (BDSC) | RRID: BDSC_51976 | 51976 |
| *CrzR-T2A-GAL4* | BDSC | RRID: BDSC_84621 | 84621 |
| *Crz<sup>10A</sup>* | A gift from Dr. Jae H. Park | (Sha, 2013) |  |
| *Crz<sup>sk2-5</sup>* | A gift from Dr. Shu Kondo | (Imura et al., 2020) |  |
| *CrzR<sup>df</sup>* | BDSC | RRID:BDSC_26828 | 26828 |
| *CrzR<sup>01</sup>* | A gift from Jae H. Park | (Sha et al., 2014) |  |
| *UAS-shi<sup>ts</sup>* | A gift from Akinao Nose | RRID:BDSC_44222 | 44222 |
| *UAS-Kir2.1* | BDSC | RRID:BDSC_6596 | 6596 |
| *UAS-TrpA1;UAS-TrpA1* | BDSC | RRID:BDSC_26263;<br>RRID:BDSC_26264 | 26263 and<br>26264 |
| *UAS-DenMark syt-GFP* | BDSC | RRID:BDSC_33064 | 33064 |
| *UAS-mCD8::GFP* | BDSC | RRID:BDSC_32186 | 32186 |
| *UAS-mCD8::RFP* | BDSC | RRID:BDSC_32219 | 32219 |
| *UAS-GCaMP6s* | BDSC | RRID:BDSC_42746 | 42746 |
| *UAS-Pink Flamingo* |  | (Kurogi et al., 2023) |  |
| *UAS-CaLexA (2-3rd)* | BDSC | RRID:BDSC_66542 | 66542 |
| *UAS-myr::GFP-T2A-RedStinger* | A gift from Toshihiro Kitamoto | (Daniels et al., 2014) |  |
| *UAS-lacZ-RNAi* | A gift from Masayuki Miura | (Kennerdell and Carthew, 2000) |  |
| *UAS-Dicer2* | BDSC | RRID:BDSC_24651 | 24651 |
| *UAS-lacZ* | BDSC | RRID:BDSC_8529 | 8529 |
| *Fru<sup>flp</sup>* | A gift from Daisuke Yamamoto | (Yu et al., 2010) |  |
| *dsx<sup>flp</sup>* | A gift from Daisuke Yamamoto | (Rezaval et al., 2014) |  |
| *UAS>stop>mCD8::GFP* | A gift from Daisuke Yamamoto | (Yu et al., 2010) |  |
| *Trhn<sup>01</sup>* | BDSC | RRID:BDSC_86147 | 86147 |
| *Trhn<sup>#21</sup>* | This study |  |  |
| *Henna<sup>#30-8</sup>* | This study |  |  |
| *UAS-Trhn-RNAi* | Vienna Drosophila Resource Center (VDRC) | RRID:Flybase_FBst0477241 | 105414 |
| *Trhn-GAL4* | A gift from Olga V. Alekseyenko | (Alekseyenko et al., 2010) |  |
| *5-HT1B<sup>AlII-V</sup>* | BDSC | RRID:BDSC_55846 | 55846 |
| *5-HT1B<sup>attP</sup>* | BDSC | RRID:BDSC_84443 | 84443 |
| *5-HT2B<sup>M107366</sup>* | BDSC | RRID:BDSC_44181 | 44181 |
| *5-HT2B<sup>attP</sup>* | BDSC | RRID:BDSC_84445 | 84445 |
| *5-HT7<sup>attP</sup>* | BDSC | RRID:BDSC_84446 | 84446 |
| *5-HT7<sup>M108096</sup>* | BDSC | RRID:BDSC_44745 | 44745 |
| *UAS-5-HT1A-RNAi<sup>SH</sup>* | VDRC | RRID:Flybase_FBst0490982 | 330198 |
| *UAS-5-HT1B-RNAi<sup>KK</sup>* | VDRC | RRID:Flybase_FBst0481570 | 109929 |
| *UAS-5-HT2A-RNAi<sup>SH</sup>* | VDRC | RRID:Flybase_FBst0490917 | 330005 |
| *UAS-5-HT2B-RNAi<sup>BL</sup>* | BDSC | RRID:BDSC_60488 | 60488 |
| *UAS-5-HT2B-RNAi<sup>KK</sup>* | VDRC | RRID:Flybase_FBst0474225 | 102356 |
| *UAS-5-HT7-RNAi<sup>SH</sup>* | VDRC | RRID:Flybase_FBst0490946 | 330023 |
| *5-HT1A-T2A-GAL4* | BDSC | RRID:BDSC_84588 | 84588 |
| *5-HT1B-T2A-GAL4* | BDSC | RRID:BDSC_84589 | 84589 |
| *5-HT2A-T2A-GAL4* | BDSC | RRID:BDSC_84590 | 84590 |
| *5-HT2B-T2A-GAL4* | BDSC | RRID:BDSC_84591 | 84591 |
| *5-HT7-T2A-GAL4* | BDSC | RRID:BDSC_84592 | 84592 |
| *SP<sup>0</sup>* | A gift from Nobuaki Tanaka | (Liu and Kubli, 2003) |  |
| *SP-promoter-SP::GFP* | Kyoto Stock Center | RRID:DGGR_116267 | 116267 |
| *dj-GFP* | BDSC | RRID:BDSC_5417 | 5417 |
| *Dup99B<sup>A2-3</sup>* | This study |  |  |
| *Dup99B<sup>attP</sup>* | BDSC | RRID:BDSC_84498 | 84498 |
| *UAS-Mip-RNAi<sup>BL</sup>* | BDSC | RRID:BDSC_41680 | 41680 |
| *Mip<sup>l</sup>* | A gift from Young-Joon Kim | (Min et al., 2016) |  |
| *Mip<sup>sk4</sup>* | A gift from Shu Kondo | (Kondo and Ueda, 2013) |  |
| *Mth<sup>attP</sup>* | BDSC | RRID:BDSC_84526 | 84526 |
| *Mth<sup>A6</sup>* | BDSC | RRID:BDSC_27904 | 27904 |
| *Prd.F-GAL4* | BDSC | RRID:BDSC_1947 | 1947 |
| *Ovu-GAL4* | A gift from Mariana Wolfner | (Chapman et al., 2003) |  |
| *DI-GAL4* | A gift from Robert Maeda | (Gligorov et al., 2013) |  |
| *Tubulin-GAL4* | BDSC | RRID:BDSC_5138 | 5138 |
| *nSyb-GAL4* | BDSC | RRID:BDSC_51941 | 51941 |
| *UAS-SPR-RNAi<sup>KK</sup>* | VDRC | RRID:Flybase_FBst0478628 | 106804 |
| *UAS-SPR-RNAi<sup>GD</sup>* | VDRC | RRID:BDSC_66872 | 7061 |
| *SPR<sup>df</sup>* | BDSC | RRID:BDSC_7708 | 7708 |
| *SPR<sup>attP</sup>* | BDSC | RRID:BDSC_84576 | 84576 |
| *SPR-T2A-GAL4* | BDSC | RRID:BDSC_84692 | 84692 |
| *SPR-T2A-LexA* | BDSC | RRID:BDSC_84431 | 84431 |
| *ChAT-GAL4, UAS-GFP* | BDSC | RRID:BDSC_6793 | 6793 |
| *13xLexAop-CD4.TdTomato (13xLexAop-mTdTomato)* | BDSC | RRID:BDSC_77139 | 77139 |
| *GMR42C06-GAL4 (R42C06-GAL4)* | BDSC | RRID:BDSC_50150 | 50150 |
| *GMR40A05-GAL4 (R40A05-GAL4)* | BDSC | RRID:BDSC_48138 | 48138 |
| *D.mel-SP-promoter-nls::GFP* | This study |  |  |
| *D.pse-SP-promoter-nls::GFP* | This study |  |  |
| *D.wil-SP-promoter-nls::GFP* | This study |  |  |
| *S.leb-SP-promoter-nls::GFP* | This study |  |  |
| *D. willistoni SPR<sup>d2</sup>* | This study |  |  |
| *D. willistoni SPR<sup>d3</sup>* | This study |  |  |
| *UAS-mSP* | A gift from Nobuaki Tanaka | (Nakayama et al., 1997) |  |

##### Wild-type strains of 12 drosophilid species

| species | From where | stock center# | Strain |
| --- | --- | --- | --- |
| *D. melanogaster* | KYORIN-Fly: Drosophila species stock center | k-s04 | (Bloomington 2057) Genome project WGS strain |
| *D. simulans* | KYORIN-Fly | E-10501 | Rakujuen (NIG) |
| *D. sechellia* | KYORIN-Fly | k-s10 | 3C |
| *D. yakuba* | KYORIN-Fly | k-s03 | Tai18E2 |
| *D. erecta* | KYORIN-Fly | k-s02 | 0224.00 |
| *D. ananassae* | KYORIN-Fly | k-s01 | AABBgl |
| *D. pseudoobscura* | KYORIN-Fly | k-s12 | MV2-25 |
| *D. persimilis* | KYORIN-Fly | k-s11 | MSH3 |
| *D. willistoni* | KYORIN-Fly | k-s13 | Gd-H4-0 |
| *D. mojavensis* | KYORIN-Fly | k-s15 | CI 12 IB-4 g8 |
| *D. virilis* | KYORIN-Fly | k-s14 | V46 |
| *S.lebanonensis* | KYORIN-Fly | E-21401 | leb0011.00 |

**Table S2.** Primer sequences for the generation of *SP*-promoter lines

Underlines indicate regions that anneal *SP*-promoter sequences. Sequences without underlines indicate the regions used for the In-Fusion reaction to ligate with the plasmids.

| promoter | Fwd<br>or<br>Rev | In-Fusion primer sequence (5'-3') | Target<br>insertion<br>length |
| --- | --- | --- | --- |
| *D. melanogaster-SP-*<br>*promoter* | Fwd | GAGCCCGGGCGAATT<u>ACGATTGGAGGAGAGTCAAC</u> | 1645 bp |
|  | Rev | GCTCCTCGAGGGATCTTTTACACCGACATTCAAGC |  |
| *D. pseudoobscura-*<br>*SP-promoter* | Fwd | GAGCCCGGGCGAATT<u>AATAGGTAAGCCGGACAAG</u> | 2061bp |
|  | Rev | GCTCCTCGAGGGATCTTCTGGTTGTCTGAGTTTTG |  |
| *D. willistoni-SP-*<br>*promoter* | Fwd | GAGCCCGGGCGAATT<u>TCAACAGGGAAAGAAAACAC</u> | 2051bp |
|  | Rev | GCTCCTCGAGGGATC<u>GTCGTTGCAGGTAAGATAAT</u> |  |
| *S. lebanonensis-SP-*<br>*promoter* | Fwd | GAGCCCGGGCGAATT<u>TTCTCCACCCATCACAAC</u> | 2188bp |
|  | Rev | GCTCCTCGAGGGATCTTGAAGGCTCTTTTCGATTG |  |

**Table S3.** gRNA sequences for mutant generation

| mutant | gRNA primer | Sequence (5'-3') |
| --- | --- | --- |
| *Trhn*<sup>#2</sup> | Trhn-gR-F1 | CTTC GGCTTCCGGGAAGAGTCTTCT |
|  | Trhn-gR-R1 | AAAC AGAAGACTCTTCCCGGAAGCC |
|  | Trhn-gR-F2 | CTTC GAAGCTCCACCTCTCACATTC |
|  | Trhn-gR-R2 | AAAC GAATGTGAGAGGTGGAGCTTC |
| *Henna*<sup>#30-8</sup> | Henna-gR-F1 | CTTC GGGATTCGGCTTACATAGTCG |
|  | Henna-gR-R1 | AAAC CGACTATGTAAGCCGAATCCC |
|  | Henna-gR-F2 | CTTC GTTCTACTCAGACGCGCAGCT |
|  | Henna-gR-R2 | AAAC AGCTGCGCGTCTGAGTAGAAC |
| *Dup99B*<sup>42-3</sup> | Dup99B-gR-F | CTTC GGATAAGGCCAATCCCAGGA |
|  | Dup99B-gR-R | AAAC TCCTGGGATTGGCCTTATCC |

| mutant | gRNA# | Sequence (5'-3') |
| --- | --- | --- |
| *D. wil-SPR* | gRNA-1(-) | ACAGTTCT\|AGAGACTGACGT TGG |
| | gRNA-2 (-) | CTACAATG\|GTTAAAATCAGC AGG |
| | gRNA-3 (+) | GAGGTGCC\|CCATCACGAGGT GGG |

**Table S4. *SP* probe sequences for in situ HCR**

|  |  |  |
| --- | --- | --- |
| *D.melanogaste*<br>*r*-SP probe1 | Probe_<br>Region<br>(5'-3') | TTCTCGTTTGCCTACTCGGCTTGGTCCAGGCCTGGGAATGGCCGTGGAA<br>TAG |
|  | sequence<br>1 (5'-3') | <u>CCTCCACGT</u><sub>aa</sub>**ACCAAGCCGAGTACGCAAACGAGAA** |
|  | sequence<br>2 (5'-3') | **CTATTCCACGGCCATTCCCAGGCCT**<sub>aa</sub><u>TCCATCTAAGCT</u> |
| *D.melanogaste*<br>*r*-SP probe2 | Probe_<br>Region<br>(5'-3') | AGCCTACAAAGTTTCCAATTCCAAGCCCCAATCCTCGTGATAAGTGGT<br>GCCG |
|  | sequence<br>1 (5'-3') | <u>CCTCCACGT</u><sub>aa</sub>**CTTGGAATTGGAACTTTGTAGGCT** |
|  | sequence<br>2 (5'-3') | **CGGCACCACTTATCACGAGGATTGG**<sub>aa</sub><u>TCCATCTAAGCT</u> |

Each probe has a split-initiator sequence [9 nucleotides (nt) or 12 nt, underlined], a linker sequence (2 nt, “aa”), and a sequence complementary to the target mRNA (25 nt, bold).

**Table S5.** RT-qPCR primers used in *Drosophila melanogaster*

| Gene names | Fwd or Rev | primer sequence (5'-3') |
| --- | --- | --- |
| *rp49* | Fwd | CGGATCGATATGCTAAGCTGT |
|  | Rev | GCGCTTGTTCGATCCGTA |
| *rp49* * | Fwd | AAGAAGCGCACCAAGCACT |
|  | Rev | TAACCGATGTTGGGCATCA |
| *SP* ** | Fwd | GAATGGCCGTGGAATAGGAA |
|  | Rev | GGCACCACTTATCACGAGGATT |
| *GFP* | Fwd | TGAACTTCAAGATCCGCCACA |
|  | Rev | GGTGCTCAGGTAGTGGTTGTC |
| *Acp63F* | Fwd | ACTGCGGAATACTTCCCGAT |
|  | Rev | CAACGCGGTTTACCATGGAG |
| *CG12828* | Fwd | AAGTGTTCTTGGTCGTGGCA |
|  | Rev | ACCGTTGATAAAGGCGCAGA |
| *CG31515* | Fwd | TTTACCTACAGTGGCTGCGG |
|  | Rev | GTCCGGTTCTGAGACGGTTT |
| *CG34130* | Fwd | GACAGCCGGCAGGATAATCA |
|  | Rev | GCTTTTGTACGAGCTCAGCG |
| *CG43145* | Fwd | TGACTGCTACGGAAGGGGTA |
|  | Rev | CCACTTGGCCATTGAGGAGT |
| *Mst57Dc* | Fwd | TAAAAACGTGGCAAAGGCCG |
|  | Rev | TTTCCAAGTCTCGACGACCG |
| *Obp56f* | Fwd | CGGCAGCTGGGGTATACAAT |
|  | Rev | TTGCTCCGAACTGATCGCAT |

\* This primer set was used in Figs. S12 and S13.

\*\* SP primer sequences are derived from Ref (Chapman et al., 2003).

**Table S6.** RT-qPCR primers used in 12 drosophilid species for Fig. 5 and Fig. S15

| Species | Fwd or Rev | Rpl32* primer sequence (5'-3') | SP#1 primer sequence (5'-3') | SP#2 primer sequence (5'-3') | SPR primer sequence (5'-3') |
| --- | --- | --- | --- | --- | --- |
| *D. melanogaster* | Fwd | AAGAAAGCGCACCAAGCACT | GAATGGCCGTGGAAATAGGAA | - | CTGCATCCGGTCTCCATGTGTC |
|  | Rev | TAACCGATGTTGGGCATCA | TTAACATCTTCCAGCCCGAGCG | - | GCAGAGTTAGCCAAACGGAGATG |
| *D. simulans* | Fwd | AAGAAAGCGCACCAAGCACT | CTCCTTGCCTACTCGGTTTGG | - | CTGCATCCGGTCTCCATGTGTC |
|  | Rev | TAACCGATGTTGGGCATCA | TTAACATCTTCCAGCCCGAGCG | - | GCAGAGTTAGCCAAACGGAGATG |
| *D. sechellia* | Fwd | AAGAAAGCGCACCAAGCACT | TGGTCTCGTTTGCCTACTCGG | - | CTGCATCCGGTCTCCATGTGTC |
|  | Rev | TAACCGATGTTGGGCATCA | CAGGCGGAGCCCAATTAAGAC | - | GCAGAGTTAGCCAAACGGAGATG |
| *D. yakuba* | Fwd | AAGAAAGCGCACCAAGCACT | CTTTGCATCGTCAGCTTGGTCC | - | CATCTCGAACTCCCTCATCTGTC |
|  | Rev | TAACCGATGTTGGGCATCA | TCTTAGCATCTTCTCCCGAGC | - | ATATACAGAGAGCCCGGTGCCG |
| *D. erecta* | Fwd | AAGAAAGCGCACCAAGCACT | GCATAGTCGGTTTGGTCCAGTC | - | CTGCATCCGGTCTCCATGTGTC |
|  | Rev | TAACCGATGTTGGGCATCA | CCCCAAATTAAGACGGCACCAC | - | GCAGAGTTAGCCAAACGGAGATG |
| *D. ananassae* | Fwd | AAGAAAGCGCACCAAGCACT | AGCATCGTCCACCCACAGAAAG | TATCCTCTAAGGCAGTGGACGC | GCAGACTAATCCCGGTTCTATAA |
|  | Rev | TAACCGATGTTGGGCATCA | GCTACTCCTTAATGCTCTGGAAG | CAAGGCCCCAGATTAAAGTCGG | TTTCAGTGACTTGCTGCCACTTAG |
| *D. pseudoobscura* | Fwd | AAGAAAGCGCACCAAGCACT | GCCGACCAAAAGTTAAGTCTAC | - | ACACCATCTCGGTGGGCTAAC |
|  | Rev | TAACCGATGTTGGGCATCA | GCTACTCCTTAATGCTCTGGAAG | - | AGTGCAGATACACCGTGCAG |
| *D. persimilis* | Fwd | AAGAAAGCGCACCAAGCACT | GCTACTCCTTAATGCTCTGGAAG | - | ACACCATCTCGGTGGGCTAAC |
|  | Rev | TAACCGATGTTGGGCATCA | GCCGACCAAAAGTTAAGTCTAC | - | AGTGCAGATACACCGTGCAG |
| *D. willistoni* | Fwd | AAGAAAGCGCACCAAGCACT | TGGCCATTGTTAGTCAGTCAATG | - | CCTGCTATGTGTACACCATTTCC |
|  | Rev | TTGAAAGCCAGTGGGCAAGCAT | ACTAGCATCTTCCACCATAGGCTG | - | TGCCATAGGTGCATGACAGACG |
| *D. mojavensis* | Fwd | AAGAAAGCGCACCAAGCACT | - | - | CCATGTGTACACCATCTCCGTC |
|  | Rev | TTGAAAGCCAGTGGGCAAGCAT | - | - | CAGCAACGCAATGTAGAAGGTGC |
| *D. virilis* | Fwd | AAGAAAGCGCACCAAGCACT | GTGCTGTTTACGTGGAGAATAC | - | ATGCACTCCGGTCTCCATGTGTC |
|  | Rev | TAACCGATGTTGGGCATCA | TGGGCTTGTCTTAGCATCTGCC | - | GCCATCGAGCATGACAGACG |
| *S. lebanonensis* | Fwd | ATCGTAAAGAAAGCGCACCAAGC | ATGCAAGCTGTGCAATCA | - | GAATCGCTGATTGTGCTCGTG |
|  | Rev | GGCCCTTAAACGACGACCAAC | TTAGCACCTGCGCACCCGAGA | - | CCAGGGGCTGGAAATATGACCG |

\* Rpl32 primer sequences except *S. lebanonensis* are derived from Tsuda et al., (2015) Genetics.
